# Lens aquaporin-5 inserts into bovine fiber cell plasma membranes through mitochondria-associated lysosome secretion

**DOI:** 10.1101/2022.02.16.480789

**Authors:** Romell B. Gletten, Lee S. Cantrell, Sujoy Bhattacharya, Kevin L. Schey

## Abstract

**PURPOSE:** To spatially map aquaporin-5 (AQP5) expression in bovine lens, molecularly characterize cytoplasmic AQP5-containing vesicles in the outer cortex, and elucidate AQP5 membrane trafficking mechanisms.

**METHODS:** Immunofluorescence was performed on bovine lens cryosections using AQP5, TOMM20, COX IV, calnexin, LC3B, LIMP-2, and connexin-50 antibodies and the fluorescent lipid membrane dye CM-DiI. AQP5 plasma membrane insertion was defined via line expression profile analysis. Transmission electron microscopy (TEM) was performed on bovine lens tissue sections to define cytoplasmic organelle identity, morphology, and subcellular localization in cortical fiber cells. Bovine lenses were treated with 10 nM bafilomycin A1 or 0.1% dimethyl sulfoxide vehicle control in *ex vivo* culture to determine changes in AQP5 plasma membrane expression.

**RESULTS:** Immunofluorescence analysis revealed cytoplasmic AQP5 expression in bovine lens epithelial cells and differentiating fiber cells. In the bovine lens cortex, complete AQP5 plasma membrane insertion occurs at *r/a* 0.951 ± 0.005. AQP5-containing cytoplasmic vesicles are spheroidal, tubular in morphology, express TOMM20, and contain LC3B and LIMP-2 as fiber cells mature. TEM analysis revealed spheroidal, tubular autophagosomes, autolysosomes, and lysosomes with degrading mitochondria. AQP5-containing cytoplasmic vesicles and autolysosomes dock and fuse with the plasma membrane. Bafiloymcin A1 treatment reduced AQP5 plasma membrane expression by 27%.

**CONCLUSIONS:** AQP5 localizes to spheroidal, tubular cytoplasmic vesicles in the differentiating bovine lens fiber cells. During fiber cell differentiation, these vesicles incorporate LC3B and fuse with LIMP-2-positive lysosomes. AQP5 trafficking to the plasma membrane occurs through lysosome secretion as a novel mechanism of AQP5 trafficking.

## Introduction

The ocular lens is a transparent, biconvex tissue in the eye that adjustably refracts visible light onto the retina enabling high acuity vision over a wide range of distances^1–3^. The optical properties of the lens are a function of its specialized cellular organization which is comprised of an anterior monolayer of epithelial cells that overlay and differentiate into lens fiber cells at the lens equator^4^. Fiber cells differentiate gradually through a process of protein synthesis, cell migration, cell elongation, and programmed organelle degradation forming a gradient of cellular differentiation and spatiotemporal protein expression from the lens periphery to the lens core.

Aquaporins (AQPs) are transmembrane water channels that represent one group of spatiotemporally expressed proteins in the lens. Mammalian lenses express AQP0^5–8^, AQP1^8,9^, and AQP5^8,10–12^. Collectively, lens AQPs regulate lens transparency and refractive index^13^ by controlling lens osmotic balance. In lens epithelial cells, AQP1 is expressed on the apical plasma membrane^14,15^ and AQP5 is expressed cytoplasmically ^15–17^. In lens fiber cells, AQP0 is expressed on the apical and basolateral plasma membranes while AQP5 is cytoplasmically expressed in the newly differentiating fiber cells and is gradually inserted into the plasma membrane as fiber cells mature ^12,15–17^.

The important role of AQP5 in maintaining lens homeostasis is implied by its expression in all mammalian lenses studied to date including human^12,18–20^, bovine^12^, mouse^11,12,16,17,19–21^, rat^12,17,21^, rabbit^8^, and dog^22^ lenses. Functional studies demonstrate the relationship between AQP5 expression and lens osmotic homeostasis. For example, basal water content and volume are increased by ∼22% and 12%, respectively, in AQP5 knockout (AQP5^*-/-*^) mouse lenses relative to wild type (AQP5^*+/+*^) lenses following osmotic perturbation in hyperglycemic media^23,24^. AQP5^*-/-*^ mouse lenses develop cataract under the same conditions but remain transparent^14,23^ following normoglycemic culture in contrast to AQP5^*+/+*^ and AQP0^*-/-*^ mice^25–29^. AQP5^*-/-*^ mice also develop age-related cataract around six months at a higher frequency than AQP5^*+/+*^ mice through upregulation of vimentin expression via miR-124–3p.1 expression^20^. The same effect is achieved in mice with a leucine to proline missense mutation at residue 51 in AQP5, AQP5^*L51P*^, which corresponds to a homologous mutation in humans (*h*AQP5^*L51P*^) associated with congenital cataract^20^.

Functional studies also demonstrate the relationship between AQP5 subcellular localization and fiber cell plasma membrane water permeability (P_H2O_) in the lens. Immunohistochemical studies show AQP5 primarily localized to the plasma membrane in the fiber cells of the mouse lens cortex and to cytoplasmic vesicles in the rat lens cortex and the extent of fiber cell permeability correlates to plasma membrane AQP5 abundance^17^. Cytoplasmic AQP5 in rat lenses is dynamically inserted into fiber cell plasma membranes in response to changes in zonular tension and thereby increases fiber cell plasma membrane water permeability (P_H2O_)^17,21^.

While the expression and functional regulation of lenticular AQP5 is currently being investigated, the molecular identity of lens AQP5-containing cytoplasmic vesicles remains unclear. AQP5 has been shown to localize to autophagosomes for degradation in both mouse and rat submandibular glands^30,31^. In this study, we spatially map AQP5 in the bovine lens, define the molecular identity of AQP5-containing cytoplasmic compartments, and investigate mechanisms of AQP5 plasma membrane insertion. In the bovine lens, we find that AQP5 is expressed cytoplasmically in the lens epithelial and outer cortical fiber cells then is gradually inserted into fiber cell plasma membranes in the inner cortex similar to other mammalian lenses. In the outer cortex, AQP5-containing cytoplasmic vesicles are identified as autolysosomes that degrade TOMM20-containing cytoplasmic vesicles possibly as a specialized method of normal lens mitochondrial autophagic degradation. These AQP5-containing autolysosomes degrade mitochondria then fuse with the plasma membrane through lysosome secretion. AQP5 plasma membrane insertion is partially inhibited by bafilomycin A1 suggesting a novel type of AQP5 trafficking through lysosome secretion, a form of type III unconventional protein secretion, in the mammalian cells.

## Materials and Methods

### Tissue

Fresh bovine lenses (1-2 years old) used for immunofluorescence were obtained from Light Hill Meats (Lynnville, TN, USA) and Cedar Hill Meat Processing (Cedar Hill, TN).

### Reagents

All chemicals, unless otherwise stated, were obtained from Sigma-Aldrich (St. Louis, MO). Bafilomycin A1 was purchased from Millipore Sigma (Burlington, MA).

### Antibodies

Affinity-purified rabbit anti-AQP5 IgG antibody targeted towards amino acid residues 249-265 of rat AQP5 (AB15858) and normal goat IgG antibody (NI02-100UG) were obtained from Millipore Sigma. Normal rabbit IgG antibodies were obtained from Cell Signaling Technology (2729; Danvers, MA). Goat anti-calnexin IgG was obtained from LifeSpan Biosciences (LS-B4403; Dallas, TX). Rabbit anti-LC3B IgG was obtained from MyBioSource, Inc. (MBS9435173; San Diego, CA). Rabbit anti-LIMP-2 IgG was obtained from Novus Biologicals, LLC (NB400-129; Centennial, CO). Mouse anti-TOMM20 IgG was obtained from Abcam (ab56783; Waltham, MA). Goat anti-connexin 50 IgG (sc-20746) and normal mouse IgG antibody were obtained from Santa Cruz Biotechnology (sc-2025; Dallas, TX). Secondary antibodies (goat anti-rabbit Alexa Fluor 488, goat anti-rabbit Alexa Fluor 647, donkey anti-rabbit Alexa Fluor 488, and donkey anti-goat Alexa Fluor 647) were obtained from Fisher Scientific. Secondary antibody used for Western blotting was obtained from Fisher Scientific (goat anti-rabbit DyLight 680, goat anti-mouse DyLight 800, and donkey anti-goat DyLight 800).

### Other Biologics

Alexa Fluor 488- or Alexa Fluor 647-conjugated wheat germ agglutinin (WGA) was used to label fiber cell plasma membranes. Vybrant™ CM-DiI Cell-Labeling Solution (cat# V22888) was used to universally label lipid membranes in fiber cells.

#### Immunofluorescence

Fresh or cultured bovine lenses were fixed in 2% paraformaldehyde-0.01% glutaraldehyde in phosphate buffered saline (PBS; 24 or 72 hours at room temperature), cryoprotected in 10% sucrose-PBS (2 days, 4°C), 20% sucrose-PBS (1 hour, room temperature), and 30% sucrose-PBS (≥ 7 days, 4°C), snap frozen in liquid nitrogen, encased in Tissue-Tek® O.C.T. Compound (Sakura Finetek USA, Inc.; Torrance, CA), cryosectioned parallel (axially) to the optic axis at 20 μm thickness using a Leica CM3050 S cryostat (Leica Biosystems Inc; Buffalo Grove, IL), and finally were transferred onto plain microscope slides. Next, lens tissue cryosections (i.e. “sections”) were triply washed in PBS, incubated in blocking solution (6% bovine serum albumin, 6% normal goat serum, in PBS) for 2-3 hours to reduce nonspecific labelling, and immunolabeled with rabbit anti-AQP5 (1:400), goat anti-calnexin (1:100), mouse anti-TOMM20 (1:200), goat anti-LC3B (1:400), or goat anti-LAMP1 (1:250) primary antibody in blocking solution (16 hours, 4°C) followed by Alexa488- or Alexa647-conjugated goat or donkey secondary antibodies in blocking solution (2 hours, room temperature). Normal rabbit IgG, normal goat IgG, and normal mouse IgG were used as host-specific negative controls for nonspecific IgG binding. In certain cases, 0.1% Triton X-100 was included in PBS washes or blocking solution prior to incubation with secondary antibodies. Following immunolabeling, sections underwent subsequent fluorescent labeling with DAPI-dilactate (1:100 in PBS) to label cellular nuclei singularly or DAPI-dilactate *in combination with* Alexa488- or Alexa647-conjugated wheat germ agglutinin (WGA; 1:100 in PBS, 1 hour, room temperature) to label fiber cell plasma membranes *or* Vybrant™ CM-DiI Cell-Labeling Solution Lipid (1:5000 in 50% ethanol-PBS) to label cellular lipid membranes. Following immunolabeling and labeling, sections were coverslipped in ProLong™Glass Antifade Mountant and imaged using a Zeiss LSM 880 confocal laser scanning microscope (Carl Zeiss Inc; White Plains, NY). Images were post-processed to improve resolution using AiryScan processing (Zeiss). Background fluorescence (i.e. fluorescence from normal IgG incubated, negative control tissue) was subtracted using Adobe PhotoShop CS6 (Adobe; San Jose, CA).

#### *Ex Vivo* Whole Lens Culture & Image Segmentation Analysis

Fresh bovine lenses were cultured in *complete medium* at 37 °C in 4% CO_2_ as outlined previously^32^ using a Forma™ Model 370 Series Steri-Cycle™ CO_2_ Incubator (ThermoFisher Scientific; Waltham, MA). Briefly, complete M199 medium consisted of M199 medium (11-150-059, Fisher Scientific), 10% fetal bovine serum, 1% penicillin, and 1% streptomycin. After 2 hours of culture, lenses free of cataract were treated with 10 nM bafilomycin A1 or 0.1% DMSO, the vehicle (negative) control, for 24 hours. Cultured bovine lenses were cryosectioned for immunofluorescence analysis and imaged as outlined above. Nikon NIS Elements version 5.3.0 software was used to perform image segmentation to quantify relative AQP5 expression changes in the cortical fiber cell plasma membranes of cultured bovine lenses due to bafilomycin A1 treatment. Relative AQP5 expression was defined as the mean intensity of AQP5 immunofluorescence with bafilomycin A1 treatment normalized to vehicle control.

#### Transmission Electron Microscopy (TEM)

Fresh bovine lenses were placed in 1% glutaraldehyde in 0.1M cacodylate buffer (2 days, room temperature). Thereafter, tissue chunks were excised from these lenses near the lens equator and incubated in 2.5% glutaraldehyde-0.1 M cacodylate (2 days, room temperature) followed by post-fixation in 1% OsO_4_ (1 hour, room temperature). The tissue was dehydrated using a graded ethanol series and infiltrated with Epon-812 (Electron Microscopy Sciences, cat# RT 13940; Hatfield, PA) using propylene oxide as the transition solvent. The Epon-812 was polymerized at 60° C for 48 hours and the samples were sectioned at 70 nm for TEM using a Leica UC7 Ultramicrotome. TEM was performed on a Tecnai T12 electron microscope (ThermoFisher Scientific) at 100 kV using an AMT CCD camera (AMT; Woburn, MA).

#### Lens Homogenization and LC-MS/MS for Identifying LIMP-2 Peptides

Frozen bovine lens anterior and posterior poles were shaved off to yield a center section approximately 3 mm thick. The cortex was removed by trephine center-punch at 7/16”. The cortex was homogenized in homogenizing buffer 25 mM Tris, 5 mM EDTA, 1 mM DTT, 150 mM NaCl, and 1 mM PMSF pH 8.0 then centrifuged at 100,000g for 30 minutes to pellet the membrane fraction. The supernatant was removed, and homogenizing buffer added before pelleting a second time to “wash” the membrane fraction. The membrane fraction was subsequently washed in homogenizing buffer with 8M urea twice then cold 0.1 M NaOH. Urea-insoluble/NaOH-insoluble membrane fraction proteins were taken up in 50 mM triethylammonium bicarbonate (TEAB) and 5% SDS and 75 μg protein was isolated. Protein isolate was reduced and alkylated with addition of 10 mM dithiothreitol and 20 mM iodoacetamide. Alkylated proteins were acidified with phosphoric acid to 2.5% and precipitated with 100 mM TEAB in methanol. Precipitate was loaded on an S-Trap micro (Protifi), washed four times with 100 mM TEAB in methanol, and digested in 0.25 μg/μL trypsin in 50 mM TEAB for 2 hours at 46°C. Peptides were eluted from the S-Trap with 66 mM TEAB, 0.2% formic acid, and then 66 mM TEAB and 50% acetonitrile (ACN). Eluted peptides were dried by a Speed Vac vacuum concentrator and rehydrated in 0.1 % triethylamine (TEA). Peptides were basic reverse phase separated on an in-house fabricated STAGE tip. Briefly, two 1.0 mm Empore C18 filter plugs were added to a pipette tip and 2 mg 5μm C18 resin (Phenomenex) was added to the top of the C18 filter plug. Peptides were loaded to the stage tip after equilibration and washed twice in 0.1% TEA. Peptides were eluted from the STAGE tip in progressive fractions of ACN (5%, 7.5%, 10%, 12.5%, 15%, 20%, 30%, 50% ACN in 0.1% TEA). Each fractions was dried separately via Speed Vac and reconstituted in 0.1% formic acid. Approximately 400 ng of each basic reverse phase fraction was separately loaded onto a trap column before separation along a 95-minute gradient from 5% - 37% ACN. Peptides were measured on a Thermo Fisher Velos Pro linear ion trap instrument operating in top15 data dependent acquisition mode. RAW files were searched with FragPipe version 17.0 and modified to accommodate the low-resolution mass analyzer used in this study. Briefly, each sample was selected as part of a single experiment and searched with MSFragger version 3.4 with precursor mass tolerance of ± 500 ppm and fragment mass tolerance of ± 0.7 Da. Peptides of length 7-50 in mass range 500-5000 with charge 1-4 were included in a database of reviewed and unreviewed proteins (downloaded 12/08/2016, length 32,167). Cysteine carbamidomethylation was included as a default modification. Up to two variable modifications of methionine oxidation and N-terminal excision were allowed per peptide. Protein level results were filtered at 5% false discovery rate (FDR) while peptides, PSMs, and ions were filtered at 1% FDR. FragPipe outputs were used for protein level data interpretation in R.

#### Statistical Analysis

All assays were conducted in at least triplicate and experimental results are represented as the data *mean* ± *standard error of the mean* (*SEM*). Statistical significance of experimental results was determined with the Student’s t-test. *P-*values ≤ 0.05 were considered statistically significant.

## Results

AQP5 has been spatially mapped in human^12^, mouse^12,15,16^, rat^12,21^, and rabbit lenses^8^. AQP5 spatial expression is broadly consistent across these species: AQP5 is cytoplasmic in lens epithelial cells and in young, differentiating fiber cells of the lens cortex and, as fiber cells mature, is gradually inserted into fiber cell plasma membranes. AQP5 expression remains localized to the plasma membrane in mature fiber cells to the lens core.

### AQP5 Localization and Membrane Insertion

Based on these findings, we hypothesized that this general AQP5 spatial expression pattern is characteristic of mammalian lenses and would also be observed in bovine lenses. Confocal microscopy analysis of bovine lens cryosections immunolabeled for AQP5 shows that AQP5 is expressed throughout the bovine lens in both lens epithelial cells and lens fiber cells (Figures 1A -1B). AQP5 expression is cytoplasmic in bovine lens epithelial cells and in the incipient fiber cells of the lens modiolus (Figures 1C-1D). AQP5 plasma membrane insertion, defined as colocalization between AQP5 immunolabeling and WGA labeling, is detectable alongside cytoplasmic AQP5 expression as differentiating cortical fiber cells mature (Figure 1E and 1F, arrowheads). AQP5 expression is completely localized to the plasma membrane in the inner cortex and remains integral to the membrane through to the lens core.

**FIGURE 1.**
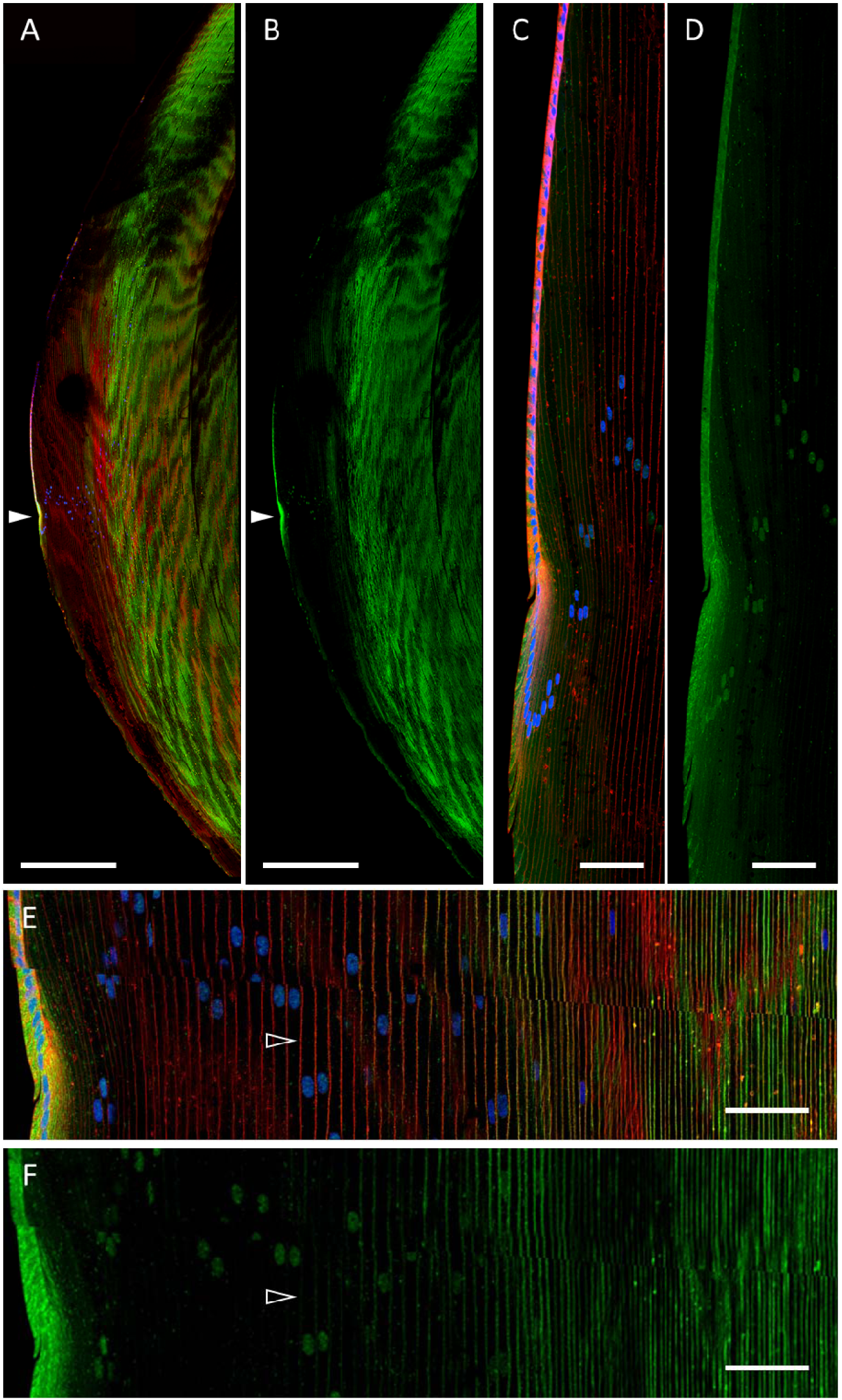
AQP5 spatial expression in the bovine lens. **A**. A low-magnification, high-resolution image of AQP5 immunolabeling (***green***) in the bovine lens with WGA labeling of plasma membranes (***red***) and DAPI labeling of cell nuclei (***blue***). The *closed arrowhead* denotes the lens modiolus. **B**. A replicate image of **A** with AQP5 immunolabeling only displayed. **C**. A medium-magnification, high-resolution image of the lens modiolus and nearby outer cortical fiber cells. **D**. A replicate image of **C** with AQP5 immunolabeling only displayed. **E**. A medium-magnification, high-resolution image of the lens cortex. The *open arrowhead* denotes the appearance of AQP5 plasma membrane insertion. **F**. A replicate image of **E** with AQP5 immunolabeling only displayed. The scale bars denote 500 μm (**A, B**) and 100 μm (**C**-**F**).

AQP5 plasma membrane insertion was quantified based on the normalized radial distance *r/a*, where *r* = *the radial distance to the lens center*, and *a* = *lens radius*^33^, as was done for previous mammalian lenses^12^. The fluorescence intensities of AQP5 and WGA in Figure 1 were converted to gray values (Figure 2A and 2B) and collinear surface plots of these gray values as a function of *r/a* quantitatively depict the spatial relationship between AQP5 expression and the plasma membrane across the lens bow region (Figure 2C-2E). AQP5 plasma membrane insertion is defined as *r/a* values with overlap between AQP5 and WGA gray values. AQP5 immunolabeling intensity peaks in the lens modiolus decreases significantly with *r/a* in the outer cortical fiber cells. The surface plot data for cytoplasmic AQP5 immunolabeling is nonuniform which reflects cytoplasmic AQP5 expression. Initial AQP5 plasma membrane insertion is detectable from *r/a* 0.979 (Figures 2C and 2D, black star) to *r/a* 0.958 (Figures 2C and 2E, blue star). Cytoplasmic AQP5 immunolabeling is detected between these *r/a* values as well. Full AQP5 membrane insertion occurs at *r/a* 0.958 (blue star), which is in the outer cortex-inner cortex transitional region. AQP5 remains localized to the plasma from 0.958 ≥ *r/a* ≥ 0.000 (data not shown).

**FIGURE 2.**
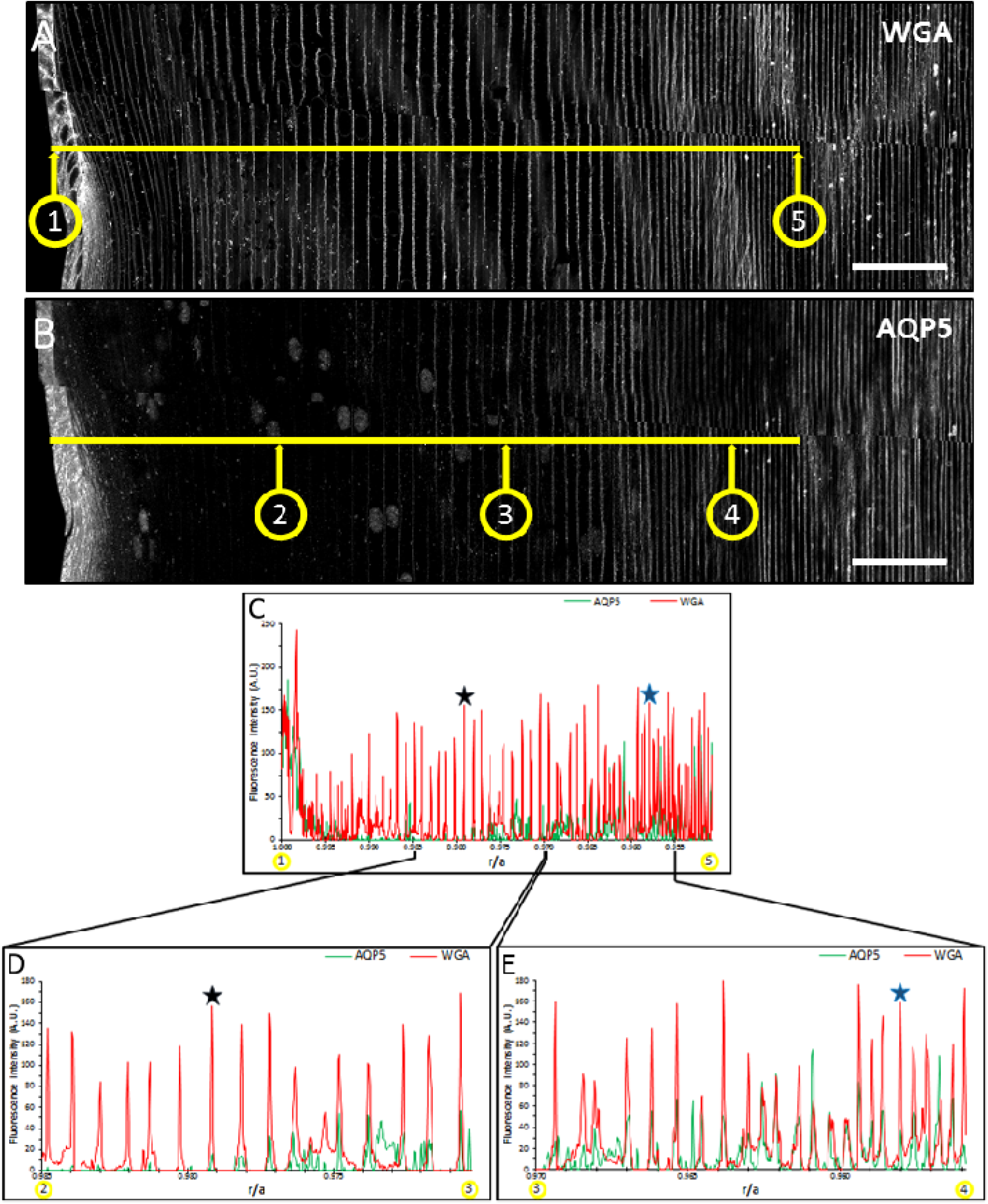
Cytoplasmic AQP5 is progressively inserted into cortical fiber plasma membranes during fiber cell differentiation in the bovine lens. **A**. A grayscale image of **Figure 1E** with WGA labeling only displayed. Distinct points in space on a lens cryosection (*n* = 1) are represented by the quotient *r/a*, where *r* = distance from any point to the center of the lens perpendicular to the optical axis, and *a* = *radius of the lens*, which is 8000 μm for this section. Point 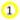 in **C** corresponds to *r/a* = 1.000 and point 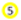 corresponds to *r/a* = 0.950. The horizontal, *yellow line* in **C** connects points 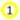 and 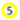. **B**. A replicate, grayscale image of **Figure 1E** with AQP5 immunolabeling only displayed. The horizontal, *yellow line* displayed is collinear to the line in **B** and connects points 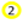, 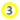, and 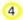 which correspond to *r/a* = 0.985, *r/a* = 0.970, and *r/a* = 0.955, respectively. **C**. Two-dimensional surface plots of the fluorescence intensity gray values of WGA in **B** (*red line*) and AQP5 in **D** (*green line*) across the *yellow line* from points 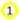 to 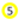. **D**. Two-dimensional surface plots of the fluorescence intensity gray values of WGA in **B** (*red line*) and AQP5 in **D** (*green line*) across the *yellow line* from points 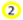 to 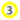. **E**. Two-dimensional surface plots of the fluorescence intensity gray values of WGA in **B** (*red line*) and AQP5 in **D** (*green line*) across the *yellow line* from points 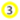 to 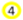. AQP5 fluorescence intensity maximizes at *r/a* = 0.998 then decreases rapidly to near undetectable levels at *r/a* = 0.989. AQP5 fluorescence intensity increases inversely with *r/a* and is partially overlapped in space by the fluorescence intensity of WGA from 0.979 (*black star*) ≥ *r/a* ≥ 0.958 (*blue star*) (**D-E**). AQP5 fluorescence intensity is entirely overlapped by that of WGA in space at 0.958 (*blue star*) (**D**). This trend continues to the lens center (0.958 ≥ *r/a* ≥ 0.000, *data not shown*). The scale bars denote 100 μm (**A**-**C**).

### Identification of AQP5 Cytoplasmic Structures

Previous studies revealed variability in the morphology of AQP5-containing cytoplasmic vesicles in mouse^12,16^, rat^12^, and human^12^ lens cortical fiber cells. To more accurately characterize AQP5-containing cytoplasmic vesicles in the bovine lens, we conducted high resolution confocal microscopy imaging of AQP5 expression in bovine lens cortical fiber cells (Figure 3). Cytoplasmic AQP5 expression in bovine lens fiber cells is localized to tubular and spheroidal cytoplasmic vesicles in the outer cortex (Figure 3, *arrowheads*). As fiber cells begin to differentiate at the lens equator, AQP5-containing cytoplasmic vesicles are less than 1 μm in diameter, up to 20 μm in length, and predominately tubular in morphology (Figure 3B, *open arrowheads*). As fiber cells mature and exit the lens modiolus, AQP5-containing cytoplasmic vesicles become predominately spheroidal and spheroidal, tubular (Figure 3, *closed arrowheads*) with spheroid regions varying up 3 μm in diameter (Figures 3C and 3D). Small puncta (< 0.45 μm) are visible that correspond to nonspecific immunofluorescence based on their appearance in negative control normal IgG immunolabeling controls (Figure 3C-3 and 3C-4).

**FIGURE 3.**
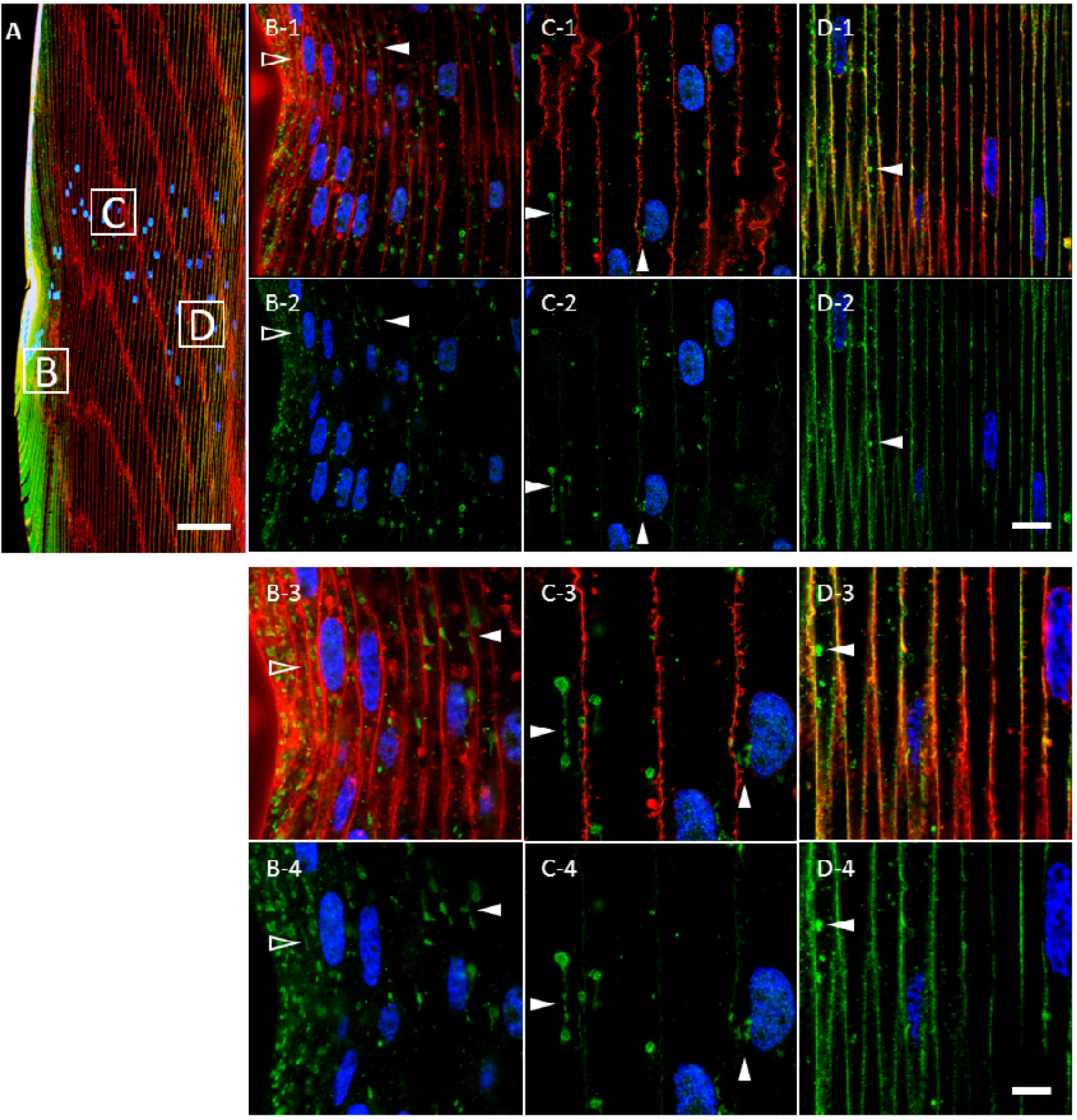
Cytoplasmic AQP5 immunolabeling in cortical fiber cells is localized to spheroidal, tubular structures in the bovine lens bow region. ***A***. A low-magnification representative image of AQP5 immunolabeling (***green***) in the bovine lens bow region. Plasma membranes (***red***) and cellular nuclei (***blue***) are labeled by WGA and DAPI, respectively. The *peripheral outer cortex* (i.e. includes *lens modiolus*), *medial outer cortex*, and *outer cortex-inner cortex transitional region* are demarcated by ***B, C***, and ***D***, respectively, and correspond spatially with the images in ***B***-***D***. ***B-1*** – ***D-1***. High-magnification images of AQP5 immunolabeling (***green***) in the cortical fiber cell regions demarcated in **A**. AQP5-containing cytoplasmic vesicles were primarily tubular in morphology in the incipient fiber cells of the lens modiolus (*open arrowheads*) and spheroidal, tubular in the cells outside of the lens modiolus (*closed arrowheads*). AQP5-containing cytoplasmic vesicles became slightly irregular in the outer cortex-inner cortex transitional region. ***B-2*** – ***D-2***. Replicate images of ***B-1*** – ***D-1*** depicting AQP5 immunolabeling and DAPI labeling only. ***B-3*** – ***D-3***. Enlarged images of AQP5-containing cytoplasmic vesicles denoted by the *arrowheads* in ***B-1*** – ***D-1***. The spheroidal and tubular domains within AQP5-containing cytoplasmic vesicles vary in size up to a maximum width and length of approximately 3 μm and 20 μm, respectively. ***B-4*** – ***D-4***. Replicate images of ***B-3*** – ***D-3*** depicting AQP5 immunolabeling and DAPI labeling only. Scale bars represent 100 μm (**A**), 10 μm (***B-1, B-2, C-1, C-2, D-1***, and ***D-2***) and 5 μm (***B-3, B-4, C-3, C-4, D-3***, and ***D-4***.

To test whether AQP5-containing cytoplasmic vesicles were morphologically distinct structures amongst cytoplasmic vesicles in bovine lens cortical fiber cells, we analyzed DiI-labeled, AQP5 immunolabeled bovine lens cryosections (Figure 4). DiI is a lipophilic dye that labels all cytoplasmic cellular lipid membranes and the plasma membrane. In the lens modiolus, tubular DiI-labeled cytoplasmic compartments overlap entirely with tubular, AQP5-containing cytoplasmic vesicles indicating these structures as morphologically unique (Figure 4A, open arrowheads). In this region, DiI-labeled spheroidal, tubular cytoplasmic compartments and AQP5-containing, cytoplasmic vesicles typically overlap, but such compartments lacking AQP5 expression are readily observable in this region (Figure 4A, striped arrowheads). In outer cortical fiber cells, with the exception of the lens modiolus, spheroidal, tubular DiI-labeled cytoplasmic compartments are identical to AQP5-containing, cytoplasmic vesicles with rare exceptions indicating that large, spheroidal, tubular cytoplasmic structures in the outer cortex are uniquely AQP5-containing cytoplasmic vesicles (Figures 4, closed white arrowheads).

**FIGURE 4.**
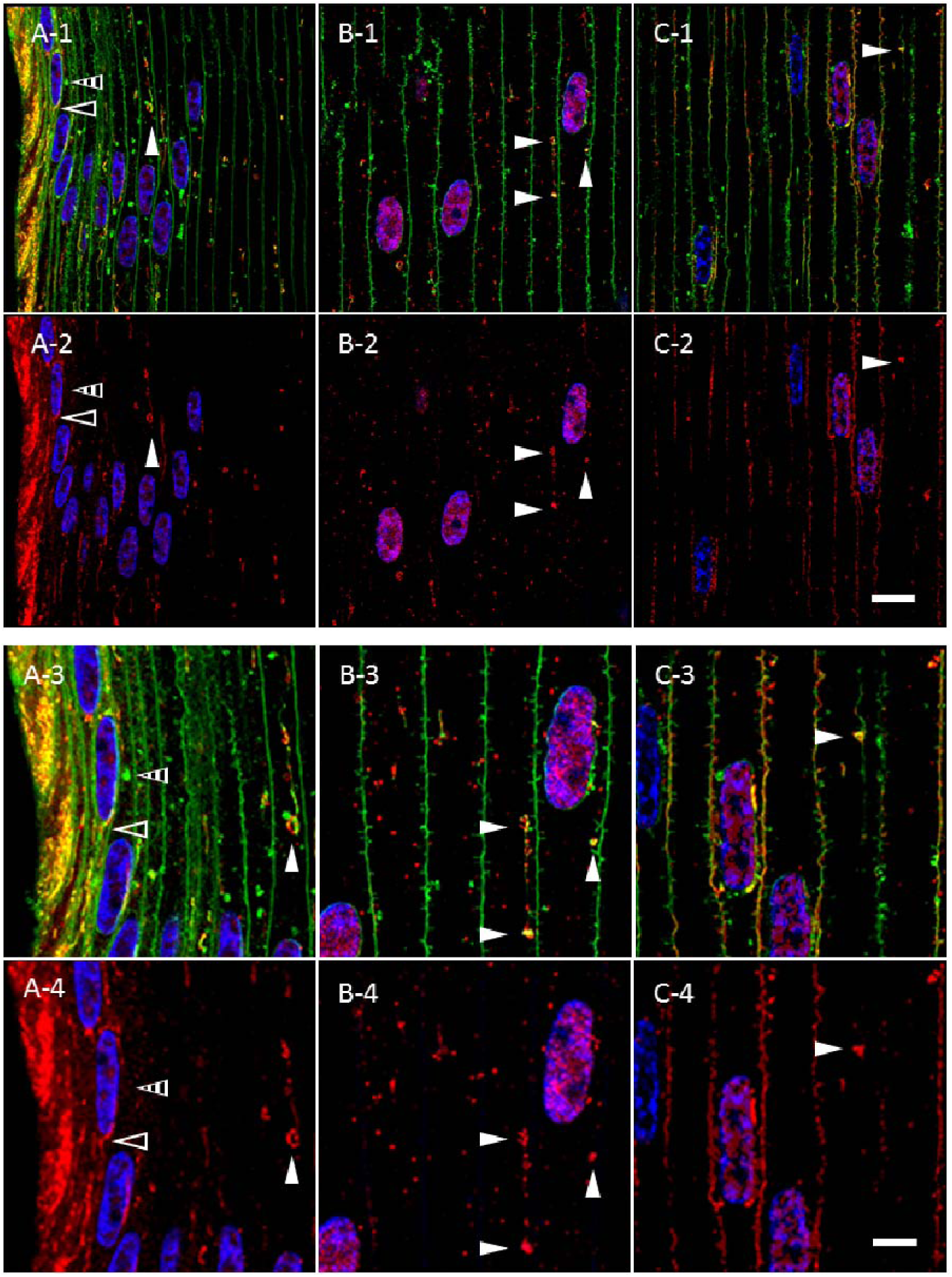
AQP5-containing, cytoplasmic vesicles represent a morphologically, distinct cluster of cytoplasmic vesicles in bovine lens cortical fiber cells outside the lens modiolus. ***A-1*** – ***C-1***. High-magnification confocal images of AQP5 immunolabeling (***red***) and DiI fluorescent labeling (***green***) in the *peripheral outer cortex* (***A***), *medial outer cortex* (***B***), and *outer cortex-inner cortex transitional region* (***C***) of the bovine lens as demarcated in ***Figure 3A***. In the lens modiolus (***A***), tubular DiI-labeled cytoplasmic compartments (*open arrowheads*) overlap with tubular, AQP5-containing cytoplasmic vesicles. In these cells, both AQP5-negative (*striped arrowheads*) and AQP5-containing (*closed white arrowheads*) DiI-labeled cytoplasmic structures are readily observable (*striped arrowheads*). In the peripheral outer cortical fiber cells outside the lens modiolus, *medial outer cortex* (***B***), and *outer cortex-inner cortex transitional region* (***C***), spheroidal, tubular DiI-labeled cytoplasmic compartments (*closed white arrowheads*) overlap with AQP5-containing, cytoplasmic vesicles with rare exceptions. ***A-2*** – ***C-2***. Replicate images of ***A-1*** – ***C-1*** with AQP5 immunolabeling and DAPI labeling only displayed. ***A-3*** – ***C-3***. Enlarged images of DiI-labeled cytoplasmic structures demarcated by arrowheads in ***A-1, B-1***, and ***C-1***. Puncta that do not colocalize with DiI meet the criteria for nonspecific immunofluorescence based on normal IgG immunolabeling negative controls. ***A-4*** – ***C-4***. Replicate images of ***A-3, B-3***, and ***C-3*** with AQP5 immunolabeling and DAPI labeling only displayed. Scale bars represent 10 μm (***A-1, A-2, B-1, B-2, C-1***, and ***C-2***) and 5 μm (***A-3, A-4, B-3, B-4, C-3***, and ***C-4***).

Immunohistochemical studies of mitochondria^34^ in chick lens fiber cells revealed structures similar in morphology to AQP5-containing cytoplasmic vesicles. To determine the molecular composition of bovine lens fiber cell AQP5-containing cytoplasmic vesicles, we tested AQP5-containing cytoplasmic vesicles for the presence of mitochondrial import receptor subunit TOM20 homolog (TOMM20) (Figure 5). TOMM20 is ubiquitously expressed in all tubular (Figure 5, *open arrowheads*) and spheroidal, tubular (Figure 5, *closed arrowheads*) AQP5-containing cytoplasmic vesicles observed. Colocalization between AQP5 and TOMM20 expression in these vesicles is high but nonuniform throughout the entirety of structures. TOMM20 expression dissipates during full AQP5 plasma membrane insertion in the bovine lens fiber cells of the outer cortex-inner cortex transition zone (Figure 5C). TOMM20 expression is not detected on the plasma membrane. The universal co-expression of AQP5 and TOMM20 within the same cytoplasmic structures in bovine lens cortical fiber cells indicates that TOMM20 is a specific molecular marker of AQP5-containing cytoplasmic vesicles. We also tested AQP5-containing cytoplasmic vesicles for the presence of cytochrome c oxidase subunit IV (COX IV; Supplementary Figure 1; S1), an inner mitochondrial membrane protein, and resident endoplasmic reticulum protein calnexin (Supplementary Figure S2; S2). COX IV expression was similar to that of TOMM20, but we observed minimal calnexin expression in AQP5-containing cytoplasmic vesicles.

**FIGURE 5.**
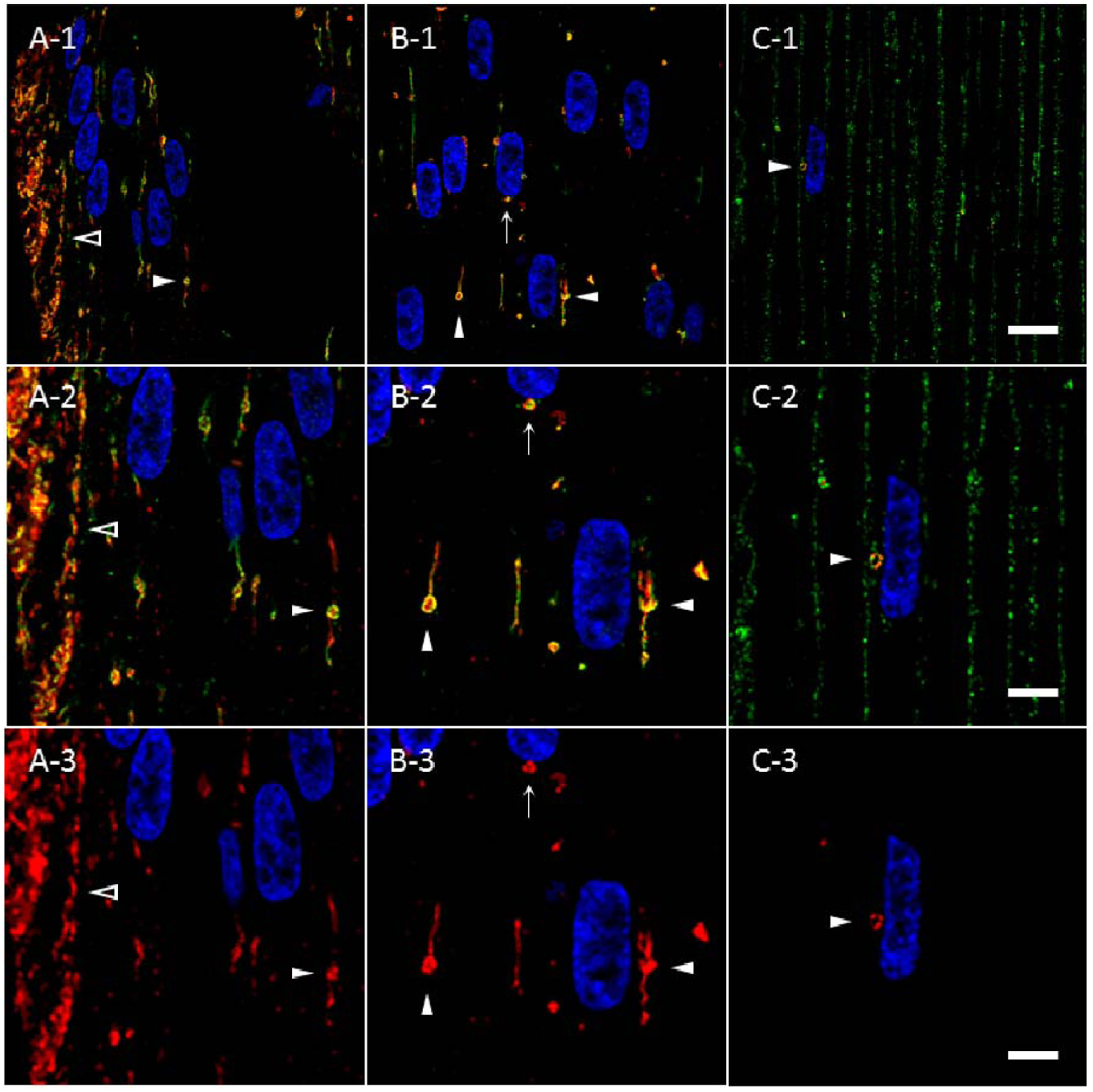
TOMM20 and AQP5 are *universally co-expressed* and *specific* molecular markers for the same cluster of cytoplasmic vesicles in the bovine lens cortex. ***A-1*** – ***C-1***. High-magnification confocal images of *AQP5* immunolabeling (***green***) and *TOMM20* immunolabeling (***red***) in the *peripheral outer cortex* (**A**), *medial outer cortex* (**B**), and *outer cortex-inner cortex transitional region* (**C**) of the bovine lens as demarcated in **Figure 3A**. Tubular (***A***, *open arrowheads*) and spheroidal, tubular (**B** and **C**; *closed arrowheads*) AQP5-containing cytoplasmic vesicles and TOMM20-containing cytoplasmic vesicles universally colocalize in the outer cortex of bovine lenses. TOMM20-containing, AQP5-containing cytoplasmic vesicles are frequently apposed to fiber cell nuclei (***5B***, *arrow*; **5C**, *closed arrowhead*). ***A-2*** – ***C-2***. Enlarged images of AQP5-containing cytoplasmic vesicles demarcated by arrowheads in ***A-1, B-1***, and ***C-1***. ***A-3*** – ***C-3***. Replicate images of ***A-2, B-2***, and ***C-2*** with TOMM20 immunolabeling and DAPI labeling only displayed. Scale bars represent 10 μm (***A-1, A-2, B-1, B-2, C-1***, and ***C-2***) and 5 μm (***A-3, B-3***, and ***C-3***).

Despite AQP5 plasma membrane insertion in the bovine lens cortex, mitochondrial degradation occurs universally within the vertebrate ocular lens^32,35–38^ and occurs through mitophagy in the mouse lens as BNIP3L/NIX expression, a mitophagy protein, is required for mitochondrial elimination to create the organelle-free zone^39^. In the mouse lens, autophagosomes are TOMM20-positive and LC3B-positive^35^. Thus, we hypothesized that AQP5- and TOMM20-containing cytoplasmic vesicles in bovine lens fiber cells might be autophagic vesicles and would also express microtubule-associated protein light chain 3B (LC3B), a specific molecular marker and peripheral membrane protein within autophagosomes and autolysosomes. We analyzed TOMM20-containing cytoplasmic vesicles in the bovine lens for LC3B expression in bovine lens fiber cells (Figure 6). In the lens modiolus, tubular TOMM20-containing cytoplasmic vesicles are readily observable as in Figure 5 (Figure 6A, *open arrowheads*), but LC3B expression in these structures is virtually undetectable. In peripheral outer cortical fiber cells outside of the lens modiolus, LC3B expression (Figure 6A, *striped arrowheads*) is sparse and sporadically localized to spheroidal, tubular TOMM20-containing cytoplasmic vesicles (Figure 6A, *closed arrowheads*). LC3B expression and colocalization within TOMM20-containing cytoplasmic vesicles increase as a function of fiber cell differentiation suggesting that mitochondria, and thereby AQP5-containing cytoplasmic vesicles, are incorporated into autophagosomes. By the outer cortex-inner cortex transitional region, TOMM20-containing cytoplasmic vesicles ubiquitously express LC3B (Figures 7B and 7C).

**FIGURE 6.**
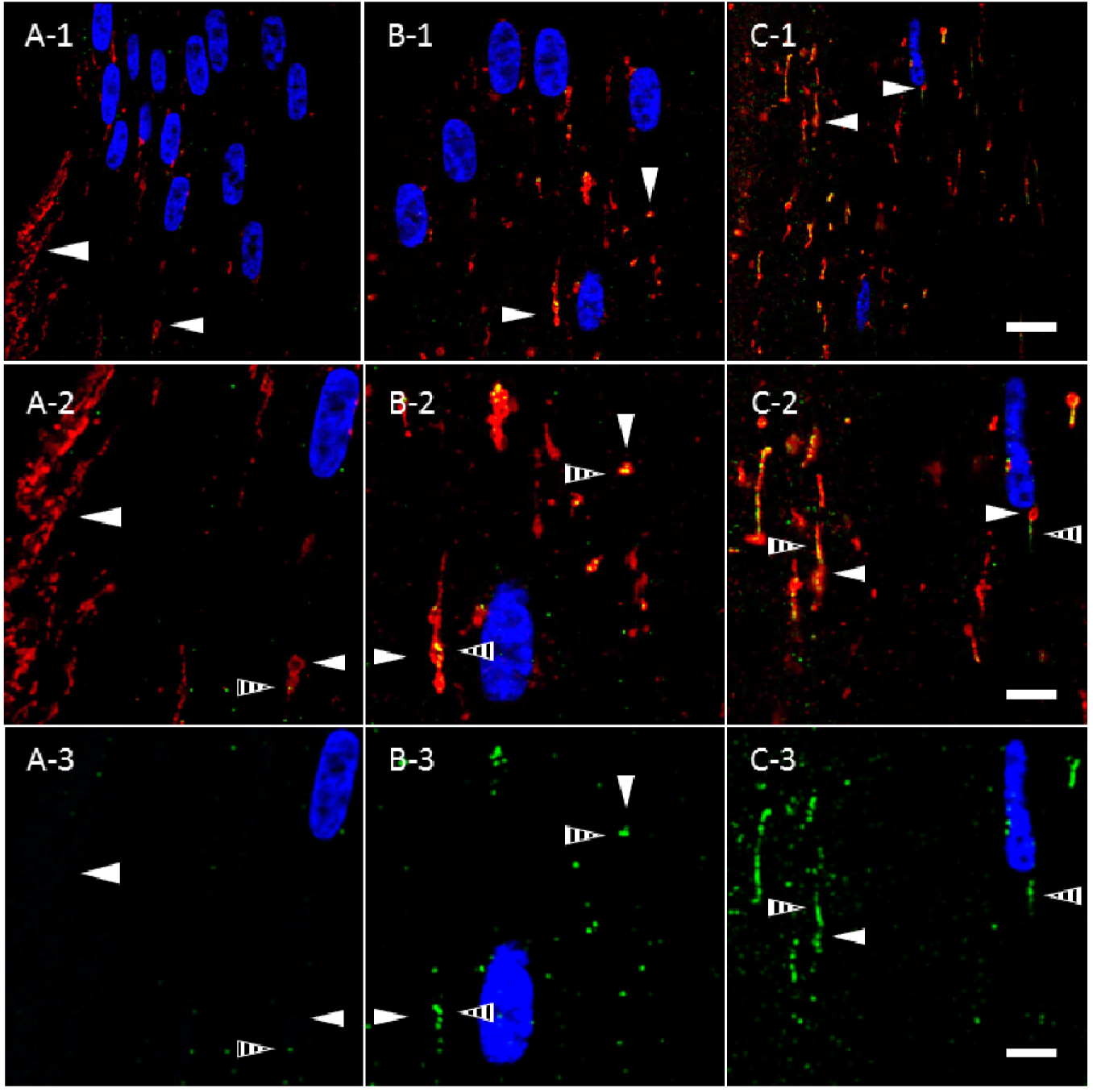
TOMM20-containing cytoplasmic vesicles express LC3B prior to complete AQP5 plasma membrane insertion in bovine lens cortical fiber cells. ***A-1*** – ***C-1***. High-magnification confocal images of *TOMM20* immunolabeling (***red***) and *LC3B* immunolabeling (***green***) in the *peripheral outer cortex* (***A***), *medial outer cortex* (***B***), and *outer cortex-inner cortex transitional region* (***C***) of the bovine lens (***Figure 3A***). In the peripheral outer cortex, LC3B is faintly expressed (***A***) in punctate, cytoplasmic vesicles (***A, B***, and ***C***; *striped arrowheads*). TOMM20-containing cytoplasmic vesicles (*closed arrowheads*) in the peripheral outer cortex colocalize minimally with LC3B. As outer cortical fiber cells mature, LC3B expression is upregulated. Thus, LC3B-containing cytoplasmic vesicles are significantly larger and more abundant in the *medial outer cortex* (***B***) and *outer cortex-inner cortex transitional region (****C***) relative to the *peripheral outer cortex*. LC3B colocalizes with the TOMM20-containing cytoplasmic vesicles in the *medial outer cortex* and the *outer cortex-inner cortex transitional region* (***B*** and ***C***). ***A-2*** – ***C-2***. Enlarged images of the TOMM20-containing cytoplasmic vesicles demarcated by the *arrowheads* in ***A-1, B-1***, and ***C-1***. ***A-3***– ***C-3***. Replicate images of ***A-2, B-2***, and ***C-2*** with LC3B immunolabeling and DAPI labeling only displayed. Scale bars represent 10 μm (***A-1, B-1***, and ***C-1***) and 5 μm (***A-2, A-3, B-2, B-3, C-2***, and ***C-3***).

**FIGURE 7.**
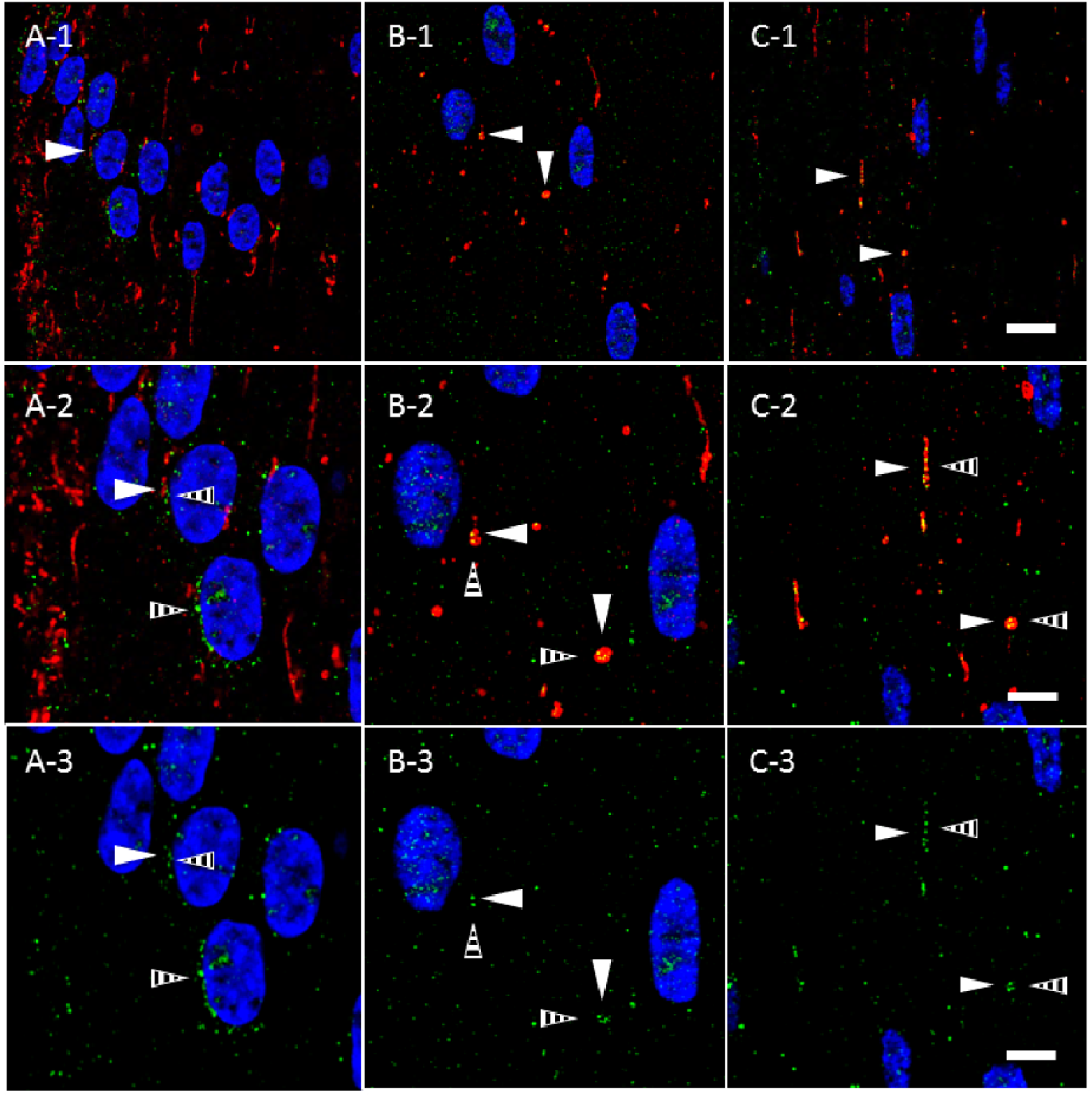
TOMM20-containing cytoplasmic vesicles express LIMP-2 prior to full AQP5 plasma membrane insertion in bovine lens cortical fiber cells. ***A-1*** – ***C-1***. High-magnification confocal images of *TOMM20* immunolabeling (***red***) and *LIMP-2* immunolabeling (***green***) in the *peripheral outer cortex* (***A***), *medial outer cortex* (***B***), and *outer cortex-inner cortex transitional region* (***C***) of the bovine lens (***Figure 3A***). In the peripheral outer cortex, LIMP-2 is faintly expressed (***A***) in cytoplasmic vesicles (***A, B***, and ***C***; *striped arrowheads*). TOMM20-containing cytoplasmic vesicles (*solid arrowheads*) in the peripheral outer cortex partially colocalize with LIMP-2. LIMP-2 expression is upregulated with fiber cell differentiation and simultaneously increased in TOMM20-containing cytoplasmic vesicles. LIMP-2 colocalizes with the TOMM20-containing cytoplasmic vesicles in the *medial outer cortex* and the *outer cortex-inner cortex transitional region* (***B*** and ***C***). ***A-2*** – ***C-2***. Enlarged images of the TOMM20-containing cytoplasmic vesicles demarcated by the *arrowheads* in ***A-1, B-1***, and ***C-1***. ***A-3*** – ***C-3***. Replicate images of ***A-2, B-2***, and ***C-2*** with LIMP-2 immunolabeling and DAPI labeling only displayed. Scale bars represent 10 μm (***A-1, B-1***, and ***C-1***) and 5 μm (***A-2, A-3, B-2, B-3, C-2***, and ***C-3***).

Autolysosomes containing mitochondria and in the process of mitophagy have been previously reported in the lens^35,38,40^. Given our observation of LC3B expression in AQP5- and TOMM20-containing cytoplasmic vesicles in bovine lens fiber cells, we hypothesized that a portion of these mitochondria would also express specific autolysosomal and lysosomal molecular markers such as lysosomal integral membrane protein 2 (LIMP-2). We analyzed TOMM20-containing cytoplasmic vesicles in bovine cortical lens fiber cells for LIMP-2 expression (Figure 7). In the lens modiolus, LIMP-2 expression in TOMM20-containing cytoplasmic vesicles is similar to that observed for LC3B. In peripheral outer cortical fiber cells outside of the lens modiolus, LIMP-2 expression (Figure 7, *striped arrows*) is also sparse and rarely overlaps TOMM20-containing cytoplasmic vesicles (Figure 7A, *closed arrowheads*). LIMP-2 expression in TOMM20-containing cytoplasmic vesicles also increases as a function of fiber cell differentiation. By the outer cortex-inner cortex transitional region, TOMM20-containing cytoplasmic vesicles express LIMP-2 with near ubiquity (Figures 7B and 7C). LIMP-2 expression in the lens was confirmed via tandem mass spectrometry (Supplementary Figure 3). These findings suggest that LC3B-positive autophagosomes merge with LIMP-2-positive lysosomes to become autolysosomes after incorporation of AQP5-containing cytoplasmic vesicles.

Given these results, we hypothesized that transmission electron microscopy (TEM) of bovine lens cortical fiber cells would reveal autophagosomes, autolysosomes, and lysosomes with similar morphology and subcellular localization to AQP5-containing cytoplasmic vesicles. TEM analysis of bovine lens cryosections revealed spheroidal, tubular AQP5-containing cytoplasmic vesicles are often in close proximity or apposed to cortical fiber cell plasma membranes (Figure 8A-1, *arrowheads*) in the bovine lens. TEM images revealed spheroidal, tubular autophagosomes and autolysosomes in close proximity (Figure 8B-1, *solid arrow*) and apposed to (Figure 8B-1, *dashed arrow*) bovine cortical fiber cell plasma membranes (Figure 8, *PM*). These structures are strikingly similar to AQP5-containing cytoplasmic vesicles in morphology. A closer look at and AQP5-containing cytoplasmic vesicle adjacent to the plasma membrane (Figure 8A-2, *open arrowhead*) reveals that the spheroid and tubular domains appear to be indistinguishable structures. In the TEM images, tubular domains in autophagosomes (Figure 8B-1, *solid arrow*), autolysosomes (Figure 8B-1, *dashed arrow*), and lysosomes (Figure 8B-2, *solid arrow*) are also at times indistinguishable from spheroid domains or separated from spheroid domains by *apparent* membranes too thin in thickness to distinguish via confocal microscopy. These results indicate that AQP5-containing cytoplasmic vesicles in the bovine lens are autophagosomes, autolysosomes, and lysosomes.

**FIGURE 8.**
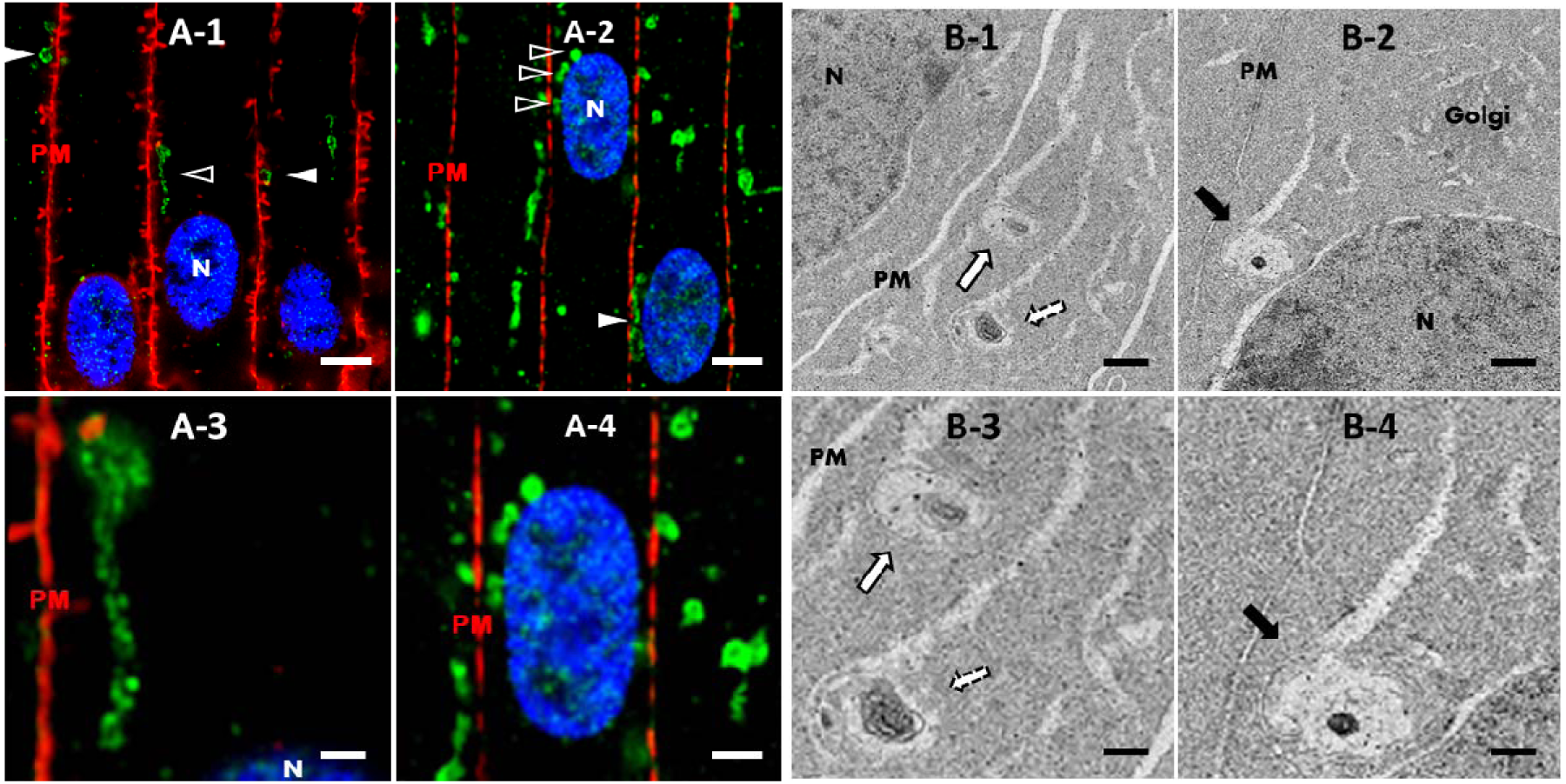
AQP5-containing cytoplasmic vesicles, autophagosomes, autolysosomes, and lysosomes in bovine lens cortical fiber cells are congruent in morphology and subcellular localization. **A**. High resolution, confocal images of AQP5 immunofluorescence (***green***) in the medial outer cortex of the bovine lens. *Plasma membranes* (**PM**) are labeled with WGA (red) (***A-1***) or connexin-50 (red) (***A-2***). *Cellular nuclei* (**N**) are labeled with DAPI staining (***blue***), respectively. AQP5-containing cytoplasmic vesicles (*arrowheads*) are distinctive amongst fiber cell cytoplasmic vesicles in the bovine lens outer cortex (***Figure 4***). AQP5-containing cytoplasmic vesicles are frequently in close proximity or apposed to fiber cell plasma membranes (***A-1***) and cellular nuclei (***A-2***). The structures demarcated with *open arrowheads* in ***A-1*** and ***A-2*** are enlarged and shown in ***A-3*** and ***A-4***, respectively. The scale bars represent 5 μm (**A-1** and **A-2**) and 2.5 μm (**A-3** and **A-4**). **B**. Transmission electron microscopy (TEM) images of and *autophagosome* (***B-1***; *solid white arrow*), and *autolysosome* (***B-1***; *dashed white arrow*), and a *lysosome* (***B-2*** *black arrow*) in outer cortical fiber cells of the bovine lens in close proximity or apposed to the plasma membrane (***B-1***) or the nucleus (***B-2***). Autophagosomes, autolysosomes, and lysosomes in these cells were roughly spheroidal and spheroidal, tubular in morphology, similar to AQP5-containing cytoplasmic vesicles. The structures demarcated with *arrows* in ***B-1*** and ***B-2*** are enlarged and shown in ***B-3*** and ***B-4***, respectively. The *Golgi apparatus* (**Golgi**) is also clearly visible. The scale bars represent 500 nm (***B-1*** and ***B-2***) and 250 nm (***B-3*** and ***B-4***).

### Mechanism of AQP5 Vesicle-Membrane Fusion

Further analysis of TEM images in the outer cortex suggested docking and fusion of autophagosomes/autolysosomes with the plasma membrane (Figure 9). We regularly observed AQP5-containing cytoplasmic vesicles apparently docked with the plasma membrane in the peripheral outer cortex prior to AQP5 plasma membrane insertion (Figure 9A, *open arrowheads*). We also observed docked vesicles undergoing the initial fusion based on colocalization between AQP5 and membrane connexin-50 immunolabeling (Figures 9A-2 and 9A-3, *arrow*). Docked AQP5-containing cytoplasmic vesicles fusing with the plasma membrane in the outer cortex-inner cortex transitional region (Figure 9A, *closed arrowhead*) were more obvious based on strong AQP5-Cx50 colocalization (Figures 9A-5 and 9A-6). In turn, we observed autolysosomal or lysosomal structures docked (Figures 9B-1 and 9B-3, *striped arrow*) and fusing (Figure 9B-2 and 9B-3, *striped arrow*) with the plasma membrane in bovine lens cortical fiber cells from the same regions.

**FIGURE 9.**
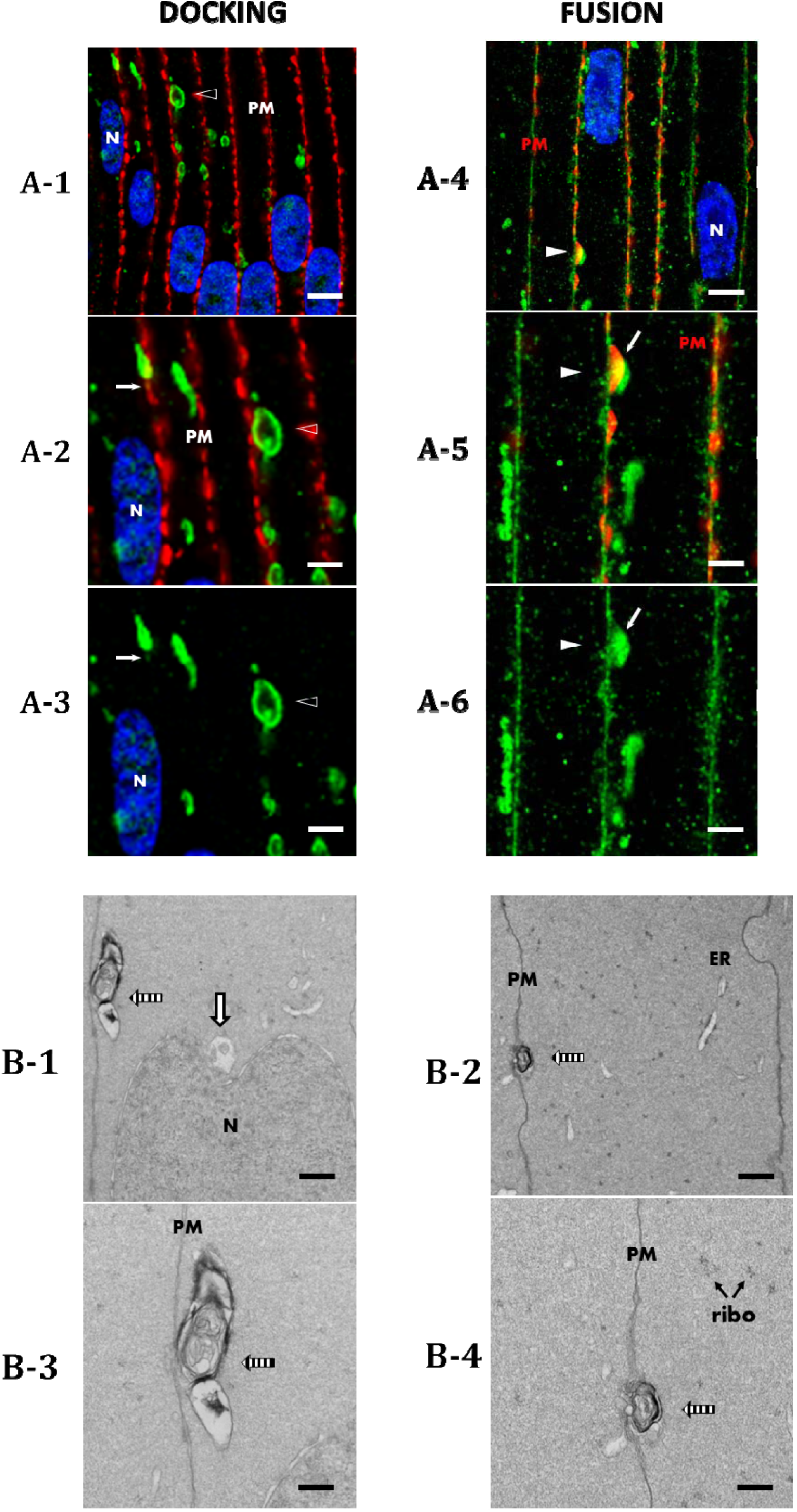
AQP5-containing cytoplasmic vesicles and autolysosomes in the bovine lens cortex dock and fuse with fiber cell plasma membranes. **A**. High resolution, confocal images of AQP5 immunofluorescence (***green***) in the *medial outer cortex* (***A-1, A-2, A-3***) and *outer cortex-inner cortex transitional region* (***A-4, A-5, A-6***) of the bovine lens. *Plasma membranes* (**PM**) are labeled with connexin-50. *Cellular nuclei* (**N**) are labeled with DAPI staining (***blue***), respectively. AQP5-containing cytoplasmic vesicles dock and fuse to fiber cell plasma membranes in the bovine lens outer cortex. AQP5-containing cytoplasmic vesicles dock by apposition to fiber cell plasma membranes (*open arrowhead*). The structures demarcated with *arrowheads* in ***A-1*** are enlarged and shown in ***A-2*** and ***A-3***. Initial fusion of a docked AQP5-containing cytoplasmic vesicle is denoted by the arrow in ***A-2*** and ***A-3***. The structures demarcated with *arrowheads* in ***A-4*** are enlarged and shown in ***A-5*** and ***A-6***. The scale bars represent 5 μm (**A-1** and **A-4**) and 2.5 μm (**A-2, A-3, A-5, A-6**). **B**. Transmission electron microscopy (TEM) images of autolysosomes (*striped arrows*) in the bovine lens outer cortex docking (***B-1*** and ***B-3***) and fusing (***B-2*** and ***B-4***) to the fiber cell plasma membrane. An autophagosome apposed to a cellular nucleus is visible as well (***B-1***, *white arrow*). The autolysosomes in ***B-1*** and ***B-3*** are enlarged and shown in ***B-2*** and ***B-4***. The *endoplasmic reticulum* (**ER**) and *ribosomes* (**ribo**) are also visible. The scale bars represent 500 nm (***B-1*** and ***B-3***) and 250 nm (***B-2*** and ***B-4***).

Together, these data support a mechanism of AQP5 plasma membrane insertion through unconventional protein secretion (UPS)^41,42^ in bovine lens cortical fiber cells – via protein targeting to autophagosomes or lysosomes predestined for fusion with the plasma membrane. These processes, “autophagosome secretion”^43–45^ and “lysosome secretion”^46–48^ respectively, are discrete mechanisms of secretion with unique molecular machineries. Secretory autophagosomes, but not degradative autophagosomes, specifically express the SNARE protein vesicle-trafficking protein SEC22β (Sec22β)^48,49^. We found that TOMM20-containing mitochondria, indistinguishable from AQP5-containing cytoplasmic vesicles through IF analysis (Figure 5), lacked Sec22β expression (Supplementary Figure 4).

The absence of Sec22β colabeling with TOMM20-containing mitochondria in combination with our TEM analyses suggests that AQP5-containing cytoplasmic vesicles become LC3B-containing autophagosomes that fuse with lysosomes that then undergo lysosome secretion in the outer cortex-inner cortex transitional region. To test this hypothesis, we examined AQP5 plasma membrane expression in *ex vivo* cultured bovine lenses treated with 10 nM bafilomycin A1 for 24 hours to inhibit autophagosome-lysosome fusion and thereby reduce lysosome secretion (Figure 10). Following *ex vivo* lens culture, AQP5 expression is still detected in the bovine lens cortex (Figure 10A-C). Relative AQP5 expression, that is mean fluorescence intensity of AQP5 with bafilomycin A1 treatment relative to vehicle control, was quantified in fiber cell plasma membranes in the bovine lens *medial outer cortex* and *outer cortex-inner cortex transitional region* (Figure 10D). Confocal microscopy imaging parameters used to acquire *ex vivo* cultured bovine lens data were kept consistent for quantitative comparison. Despite sample variation, there was no significant difference in relative AQP5 expression in fiber cell plasma membranes of the *medial outer cortex*. In contrast, relative AQP5 fiber cell plasma membrane expression in the *outer cortex-inner cortex transitional region*, where full insertion of AQP5-containing cytoplasmic vesicles with fiber cell plasma membrane occurs, was decreased by approximately 27%. These results are consistent with our hypothesis of AQP5 lysosome secretion in the bovine lens.

**FIGURE 10.**
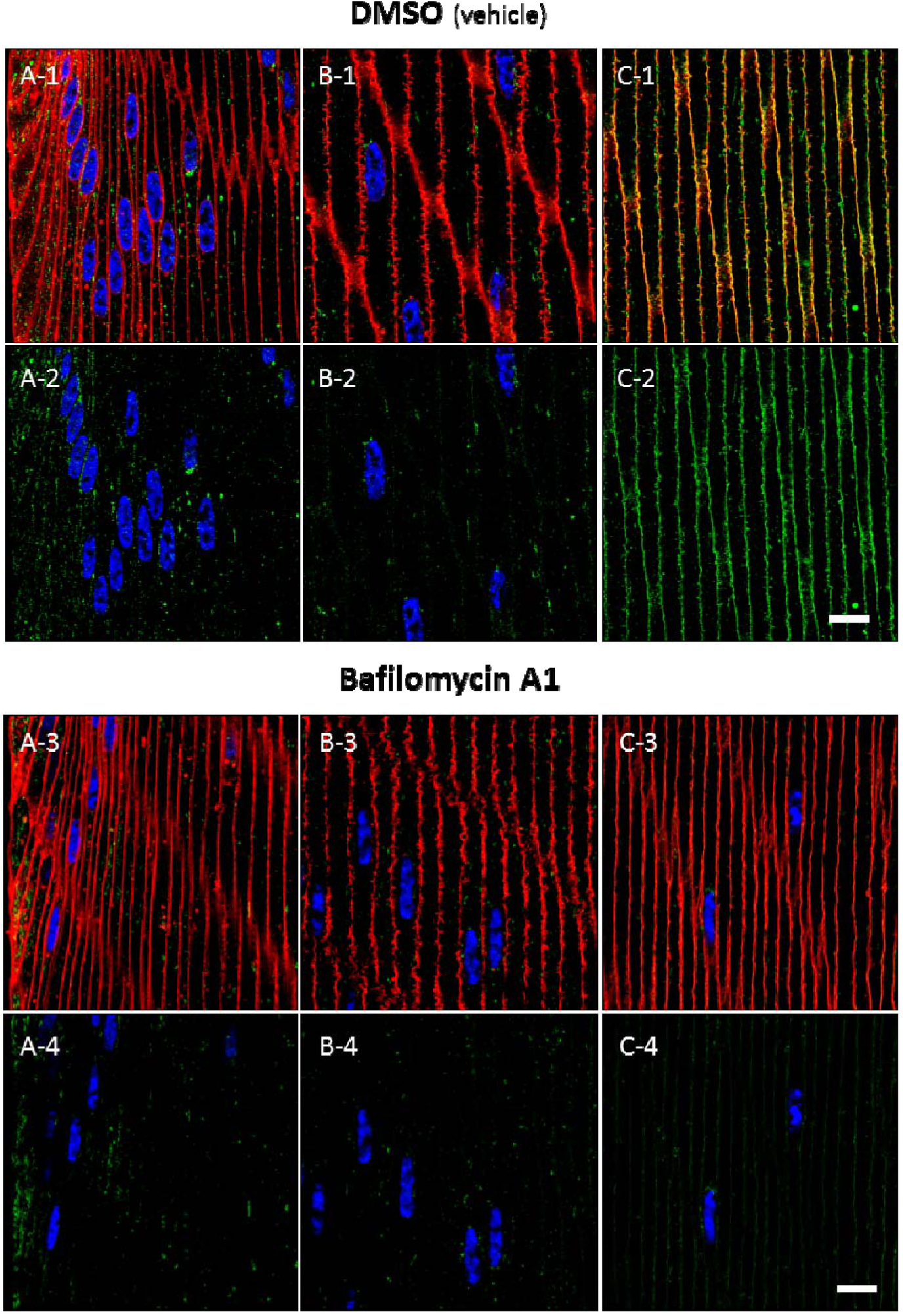
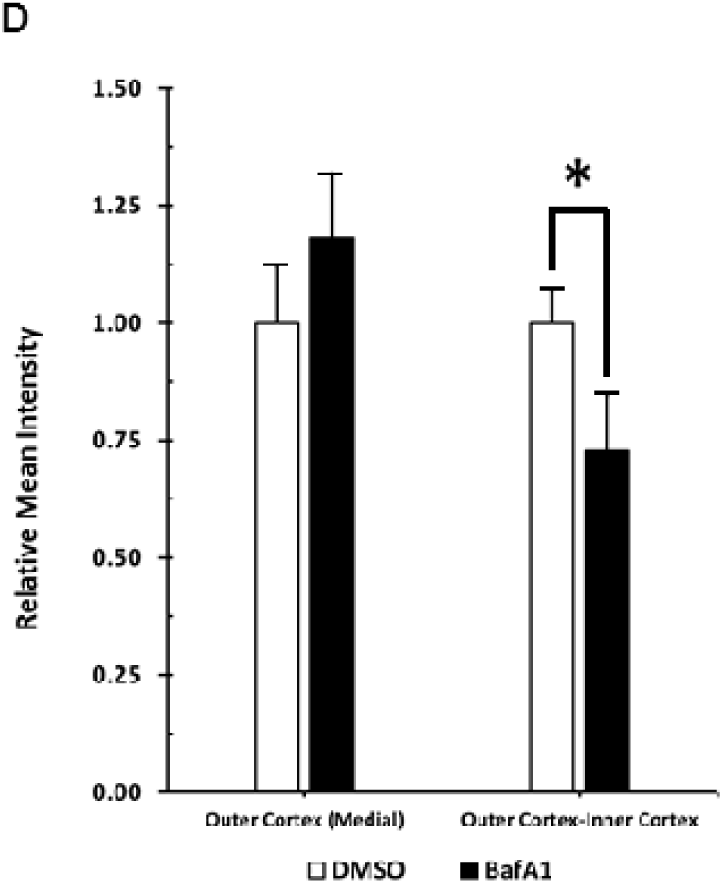
Autophagosome-lysosome fusion inhibition via bafilomycin A1 treatment decreases AQP5 plasma membrane expression in the bovine lens cortex. ***A-1*** – ***C-1***. High-magnification confocal images of *AQP5* immunolabeling (***green***) and *WGA* labeling (***red***) in the *peripheral outer cortex* (***A***), *medial outer cortex* (***B***), and *outer cortex-inner cortex transitional region* (***C***) of the bovine lens following 24 hours *ex vivo* culture in complete M199 medium with vehicle (0.1% DMSO) (***Figure 3A***). ***A-2*** – ***C-2***. Replicate images of ***A-1, B-1***, and ***C-1*** with AQP5 immunolabeling and DAPI labeling only displayed. ***A-3*** – ***C-3***. High-magnification confocal images of *AQP5* immunolabeling (***green***) and *WGA* labeling (***red***) in the *peripheral outer cortex* (***A***), *medial outer cortex* (***B***), and *outer cortex-inner cortex transitional region* (***C***) of the bovine lens following 24 hours *ex vivo* culture with 10 nM bafilomycin A1 (***Figure 3A***). ***A-4*** – ***C-4***. Replicate images of ***A-3, B-3***, and ***C-3*** with AQP5 immunolabeling and DAPI labeling only displayed. Scale bars represent 10 μm (***A, B***, and ***C***) ***D***. Quantification and statistical analysis of the relative AQP5 expression, defined as relative mean intensity, in fiber cells of the *medial outer cortex* and *outer cortex-inner cortex transitional region* of bovine lens cryosections following 24 hours *ex vivo* culture with vehicle control 0.1% DMSO (DMSO, n = 7) or with 10 nM bafilomycin A1 (BafA1, n = 8). Two-tailed Student’s t-test was significant (*) a *p*-value of 0.032.

## Discussion

The goals of this study were to determine how bovine lenticular AQP5 expression patterns compare to other mammalian lenses, to determine the subcellular localization of AQP5-containing cytoplasmic vesicles, and to identify potential trafficking mechanisms of AQP5-containing cytoplasmic vesicles to the plasma membrane. In this study, we reveal that bovine lens AQP5 expression patterns are similar to that of other mammalian lenses, being cytoplasmic in lens epithelial cells and young, differentiating lens fiber then shifting to lens fiber cell plasma membranes with cellular maturation. We have also for the first time defined AQP5 subcellular localization in ocular lens cortical fiber cells in detail and discovered that cytoplasmic AQP5 trafficks to the plasma membrane in these cells via lysosome secretion, a novel mechanism of aquaporin trafficking.

AQP5 spatial expression has previously been investigated in mouse^12,15–17^, rat^12,17^, rabbit^8^, and human^12^ lenses. While Grey et al., 2013 validated AQP5 expression in bovine lenses via Western blot analysis^12^, this study represents the first histological study of AQP5 spatial expression in the bovine lens. AQP5 is ubiquitously expressed throughout the bovine lens; being cytoplasmic in epithelial and fiber cells and gradually trafficking to plasma membrane during fiber cell differentiation. This spatial expression pattern is consistent with previously studied lenses and further suggests that this general expression pattern is characteristic of AQP5 expression in all mammalian lenses. In bovine lenses we measured AQP5 plasma membrane insertion to occur at a normalized r/a value of 0.958. Grey et al 2013 defined this value in mouse and rat lenses as *r/a* 0.95 and *r/a*∼ 0.75 to ∼0.65, respectively^12^. This equivalence suggests similarity in regulation of AQP5 plasma membrane insertion in mouse and bovine lenses relative to rat lenses, but the implications of these values remain to be investigated.

This is also the first study to characterize AQP5-containing cytoplasmic vesicles in a mammalian lens for intrinsic properties such as morphology or molecular composition. The presence of micrometer-scale tubular and spheroidal, tubular AQP5-containing cytoplasmic compartments (Figure 3) is a novel finding in cortical lens fiber cells. Within bovine cortical lens fiber cells, AQP5-containing cytoplasmic vesicles are also morphologically distinct (Figures 4A-3 and 4A-4) which enables their identification via downstream TEM analysis.

Molecular analysis revealed that AQP5-containing cytoplasmic vesicles and mitochondria are indistinguishable cytoplasmic compartments in bovine lens cortical fiber cells. TOMM20 (Figure 5), a protein subunit within the translocase of the outer mitochondrial membrane, and cytochrome c oxidase IV (COX IV) (Figure S1), a mitochondrial marker protein in the electron transport chain, both colocalize to AQP5-containing cytoplasmic vesicles in bovine lens cortical fiber cells, indicating that AQP5 is either incorporated into mitochondria or into mitochondria-containing cytoplasmic vesicles. TEM analysis in bovine lens cortical fiber cells revealed widespread incorporation of mitochondria into autophagosomes and autolysosomes (Figure 8B), demonstrating that AQP5-containing cytoplasmic vesicles are likely incorporated into autophagosomes and then into autolysosomes. These data are consistent with TEM analysis of fiber cells in human and embryonic chick lenses conducted by Costello et al, 2013 in which mitochondria are similarly incorporated into autolysosomes^35^. Calnexin IF analysis suggested that AQP5-containing cytoplasmic vesicles are not endoplasmic reticular compartments (Figure S4).

Mitochondria undergo autophagic degradation to generate an organelle-free zone (OFZ) in the lens to prevent light scatter^35^. TOMM20 colocalizes with LC3B, a nucleation and specific protein marker of autophagosomes, in the embryonic chick lens prior to mitochondrial degradation. IF analysis revealed LC3B colocalization with TOMM20 in bovine lens cortical fiber cells (Figure 6) that peaks within the outer cortex-inner cortex transitional region. LIMP-2 is a β-glucocerebrosidase receptor and specific protein marker of lysosomes^50^ and colocalization of LIMP-2 with TOMM20 (Figure 7), similar to LC3B, indicates lysosomal involvement in mitochondrial degradation. This finding is in agreement with previous data that show LC3B expression peaks and autolysosomes are abundant in the outer cortex-inner cortex transitional region in mice just prior to mitochondrial elimination to form the OFZ^37,38,51^.TOMM20 and LC3B immunolabeling fade to become undetectable in mature fiber cells. AQP5 also is fully inserted into bovine lens fiber cell plasma membranes the outer cortex-inner cortex transitional region. The disappearance of TOMM20 and LC3B immunolabeling along with our TEM data indicate that mitochondria are incorporated into autophagosomes that merge with lysosomes to become autolysosomes and degrade mitochondria, consistent with the conservation of mitochondrial autophagy in the lens across species. Genetic deletion studies suggest that lenticular mitochondrial autophagy is specific and technically mitophagy in the mouse lens^39^. This possibility remains to be tested in the bovine lens.

The persistence of AQP5 immunolabeling despite mitochondrial degradation and AQP5 plasma membrane insertion implies that AQP5-containing cytoplasmic vesicles are trafficked to the plasma membrane through the unconventional protein secretion pathway (UPS) of lysosome secretion^46^. IF and TEM analysis show that AQP5-containing cytoplasmic structures and autolysosomes dock and fuse with fiber cell plasma membranes (Figure 9). This is the first study to report lysosome secretion as trafficking mechanism of plasma membrane insertion for AQP5, for aquaporins as a protein family in any tissue, and for any protein in lens fiber cells. The absence of Sec22β colocalization with TOMM20 (Figure S2) indicates that mitochondria, and therefore AQP5, are not incorporated into secretory autophagosomes and that AQP5-containing cytoplasmic vesicles undergo lysosome secretion. Autophagosome secretion and lysosome secretion are classified as Type III unconventional protein secretion mechanisms^42,48^. Thus, while mitochondrial degradation through autophagy in bovine lens cortical fiber cells is consistent with other mammalian lenses, our data suggests that AQP5 is uniquely trafficked in bovine lenses through secretory lysosomes via LC3B-containing autophagosomes. Results from bafilomycin A1 treatment of *ex vivo* lens cultured bovine lenses support this hypothesis (Figure 10). Upon treatment, relative AQP5 fiber cell plasma membrane expression in bovine lenses decreases by approximately 27% in the *outer cortex-inner cortex transitional region*, where AQP5-containing cytoplasmic vesicles primarily traffick to the plasma membrane (Figure 10D). This is consistent with bafilomycin A1 inhibition of autophagosome-lysosome fusion and suggests an accumulation of AQP5-containing autophagosomes unable to fuse with lysosomes for secretion.

Results from bafilomycin A1 treatment also provide further evidence that AQP5 functions as a regulatory water channel whose trafficking is inducible in the mammalian lens. Petrova et al has previously demonstrated dynamic AQP5 subcellular localization changes in response to changes in zonular tension via mechanosensitive TRPV1 channel agonism^17,21^. Since AQP5 plasma membrane insertion significantly increases fiber cell water permeability in both mouse and rat lenses^17^, regulation of AQP5 subcellular localization can alter lens water homeostasis. Whether regulation of AQP5 subcellular localization via TRPV1 channels involves lysosome secretion remains to be investigated.

In non-lenticular tissues, cytoplasmic AQP5 undergoes regulated secretion to the plasma membrane in the salivary glands and in model cell systems upon M_3_ muscarinic acetylcholine receptor (AChR) activation^52,53^, β-adrenergic receptor activation^15,54,55^, TRPV4 activation^56,57^, and osmotic perturbation^56,57^. Downstream signal transduction effectors of these activators such as nitric oxide, protein kinase G, protein kinase A (PKA) activity, and calcium have all been implicated in the alteration of AQP5 plasma membrane trafficking. To our knowledge, this is the first study to characterize the subcellular localization of cytoplasmic AQP5 vesicles targeted to the plasma membrane for regulated secretion. Our results demonstrate a novel mechanism of cytoplasmic AQP5 trafficking to the plasma membrane. The effects of nonlenticular effector proteins such as the β-adrenergic receptors or corresponding downstream effectors such as PKA on AQP5 plasma membrane trafficking in the lens remain to be investigated.

In addition to the novel trafficking mechanism elucidated in this study, our results demonstrate possible AQP5 expression in viable mitochondria, which, if confirmed, along with aquaporin-8 (AQP8)^58^ would represent the second aquaporin discovered in mitochondria. These results raise a variety of questions about the conservation of this AQP5 lysosome secretion across mammalian lenses and the implications of AQP5 in active mitochondria. Future studies should address these and other relevant exciting questions both in the mammalian lens and in other tissues expressing AQP5.

## Conclusions

In summary, bovine lenticular AQP5 expression patterns are similar to AQP5 expression patterns in other mammalian lenses being cytoplasmic in the epithelial cells and young fiber cells then gradually trafficking to the plasma the plasma membrane during fiber cell differentiation. In the outer cortical bovine lens fiber cells, cytoplasmically expressed AQP5 is localized to LC3B-positive autophagosomes that merge with lysosomes to become LIMP-2-positive autolysosomes. These autolysosomes degrade mitochondria in differentiating lens fiber cells and fuse with fiber cell plasma membranes through lysosome secretion, resulting in plasma membrane insertion of AQP5. This trafficking mechanism is bafilomycin A1-sensitive indicating the importance of autophagosome-lysosome fusion for AQP5 plasma membrane insertion in the bovine lens.

## Acknowledgements

This work was supported by NIH grants R01 EY013462 and P30 EY008126.

