## Supplementary Data for "Lens aquaporin-5 inserts into bovine fiber cell plasma membranes through mitochondria-associated lysosome secretion"

Romell B. Gletten[1], Lee S. Cantrell[1], Sujoy Bhattacharya[2], and Kevin L. Schey[1]
[1] Department of Biochemistry, Vanderbilt University, Nashville, Tennessee, United States
[2]Department of Ophthalmology and Visual Sciences

Correspondence: Kevin L. Schey, Mass Spectrometry Research Center, 465 21st Avenue South, Suite 9160, Medical Research Building III, Vanderbilt University, Nashville, TN 37232-8575, USA


### SUPPLEMENTARY DATA

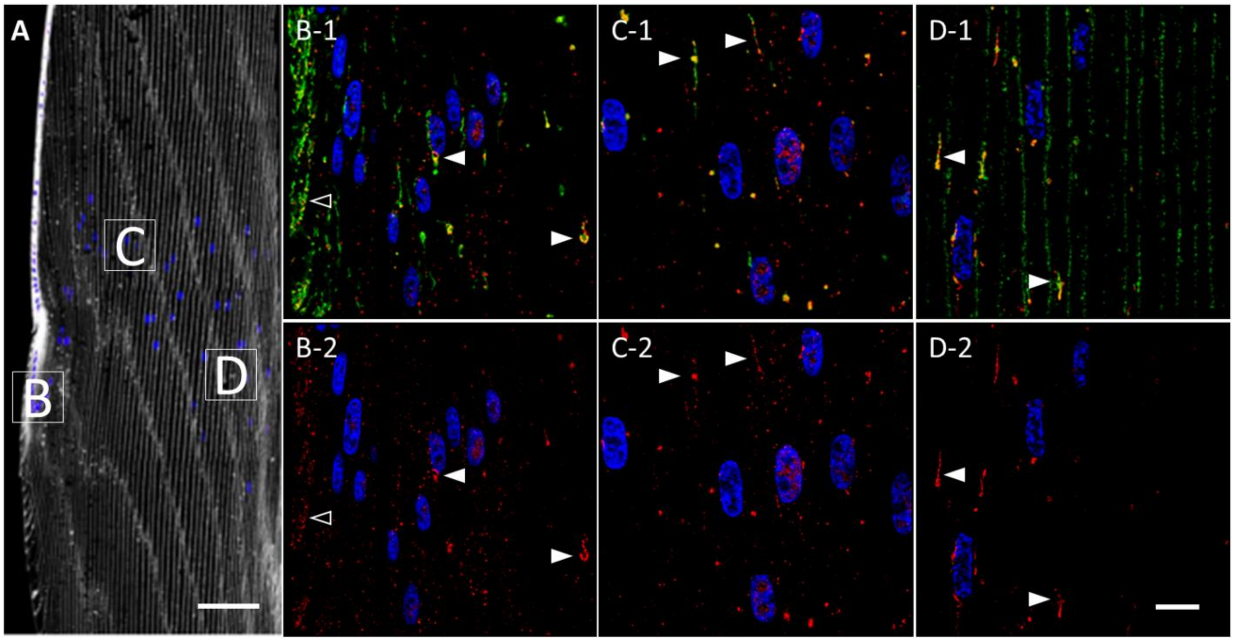


**S1. Cytochrome c oxidase subunit IV (COX IV) and AQP5 are *universally co-expressed* and *specific* molecular markers for the same cluster of cytoplasmic vesicles in the bovine lens cortex.**

***A*.** A low-magnification representative image of the bovine lens bow region with plasma membranes (***white***) and cellular nuclei (***blue***) labeled by WGA and DAPI, respectively. The *peripheral outer cortex* (i.e. includes *lens modiolus*), *medial outer cortex*, and *outer cortex-inner cortex transitional region* are demarcated by ***B***, ***C***, and ***D***, respectively, and correspond spatially with the images in ***B***-***D***.

***B-1* – *D-1*.** High-magnification confocal images of *AQP5* immunolabeling (***green***) and *cytochrome c oxidase subunit IV* (*COX IV*) immunolabeling (***red***) in the *peripheral outer cortex* (**A**), *medial outer cortex*

(**B**), and *outer cortex-inner cortex transitional region* (**C**) of the bovine lens as demarcated in **Figure S1A**.

Tubular (***A***, *open arrowheads*) and spheroidal, tubular (**B** and **C**; *closed arrowheads*) AQP5-containing cytoplasmic vesicles and COX IV-containing mitochondria universally colocalize in the outer cortex of bovine lenses.

***B-2*** – ***D-2*.** Replicate images of ***A-1***, ***B-1***, and ***C-1*** with COX IV immunolabeling and DAPI labeling only displayed.

Scale bars represent 100 µm (***A***) and 10 µm (***B***, ***C***, and ***D***).

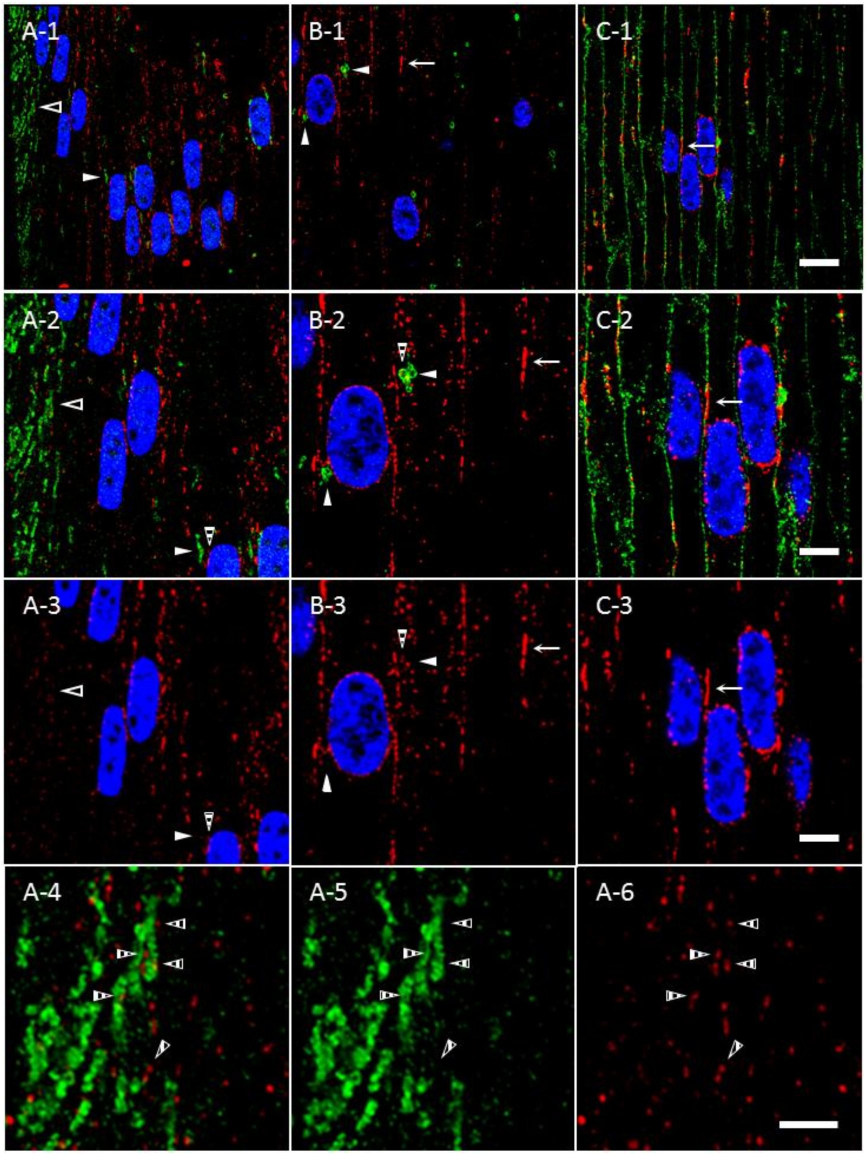


**S2. AQP5-containing cytoplasmic vesicles and calnexin-containing endoplasmic reticula fail to overlap in the bovine lens cortex.**

***A-1*** – ***C-1*.** High-magnification confocal images of *AQP5* immunolabeling (***green***) and *calnexin* labeling (***red***) in the *peripheral outer cortex* (***A-1***), *medial outer cortex* (***B-1***), and *outer cortex-inner cortex transitional region* (***C-1***) of the bovine lens (demarcated in **Figure 3A**). Calnexin is an integral membrane protein in the endoplasmic reticulum (ER). In the *peripheral outer cortex* and *medial outer cortex*, calnexin is cytoplasmically expressed in irregular, punctate vesicles or tubular, plaque-like ER compartments apposed to fiber cell plasma membranes. Calnexin is also expressed in the perinuclear ER (***B-1***, ***B-2***, ***C-1***, and ***C-2***). By the outer cortex-inner cortex transitional region, calnexin expression is primarily localized to the perinuclear ER and tubular ER apposed to fiber cell plasma membranes (***B*** and ***C***, arrows).

AQP5-containing cytoplasmic vesicles lack significant calnexin expression. However, calnexin-containing ER form multiple appositions (***A-3***, ***A-4***, ***A-5***, ***A-6***, ***B-3***, and ***B-4,*** *striped arrowheads*) with tubular (***A***, *open arrowheads*) and spheroidal, tubular (***A*** and ***B***; *closed arrowheads*) AQP5-containing cytoplasmic vesicles.

***A-2*** – ***C-2*****.** Enlarged images of the AQP5-containing cytoplasmic vesicles demarcated by the *arrowheads* in ***A-1*** and ***B-1*** and of the calnexin-containing tubular ER compartment demarcated by the *arrows* in ***B-1*** and ***C-1***.

***A-3*** – ***C-3*****.** Replicate images of ***A-2***, ***B-2***, and ***C-2*** with calnexin immunolabeling and DAPI labeling only displayed.

***A-4*****.** Enlarged image of tubular AQP5-containing cytoplasmic vesicles demarcated by the *open arrowhead* in ***A-2***. These vesicles typically form several appositions with calnexin-containing ER (*striped arrowheads*)

***A-5*****.** A replicate image of ***A-4*** with AQP5 immunolabeling only displayed.

***A-6*****.** A replicate image of ***A-4*** with calnexin immunolabeling only displayed.

Scale bars represent 10 µm (***A-1***, ***B-1***, and ***C-1***) and 5 µm (***A-2***, ***A-3***, ***B-2***, ***B-3***, ***C-2***, and ***C-3***).

***A.***

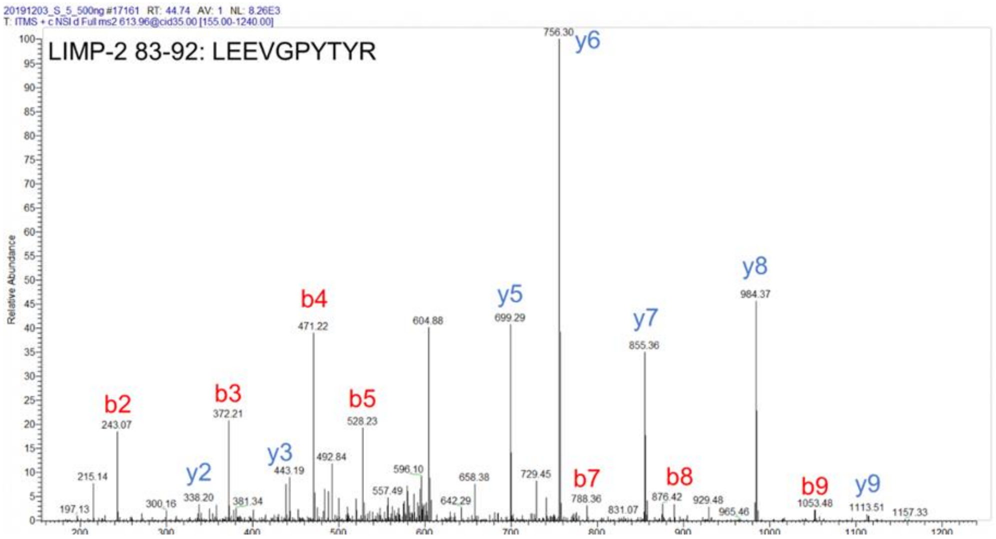

***B.***

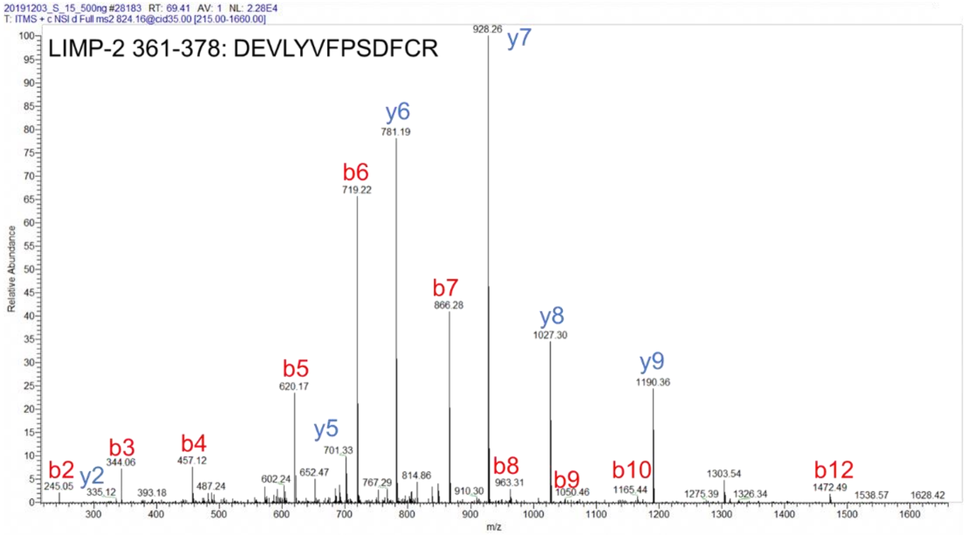


**S3. Tandem mass spectra identifying LIMP-2 in bovine lens cortical fiber cells.**

***A.*** High mass resolution CID spectrum of doubly-charged LIMP-2 peptide 82-93, m/z 824.16, acquired on a Thermo Fisher Velos Pro linear ion trap. The observed mass corresponds to one carbamidomethylated cysteine residue on the peptide (b12, y2). The peptide amino acid sequence is included above.

***B.*** High mass resolution CID spectrum of doubly-charged LIMP-2 peptide 361-378, m/z 824.16, acquired on a Thermo Fisher Velos Pro linear ion trap. The observed mass corresponds to one carbamidomethylated cysteine residue on the peptide (b12, y2). The peptide amino acid sequence is included above.

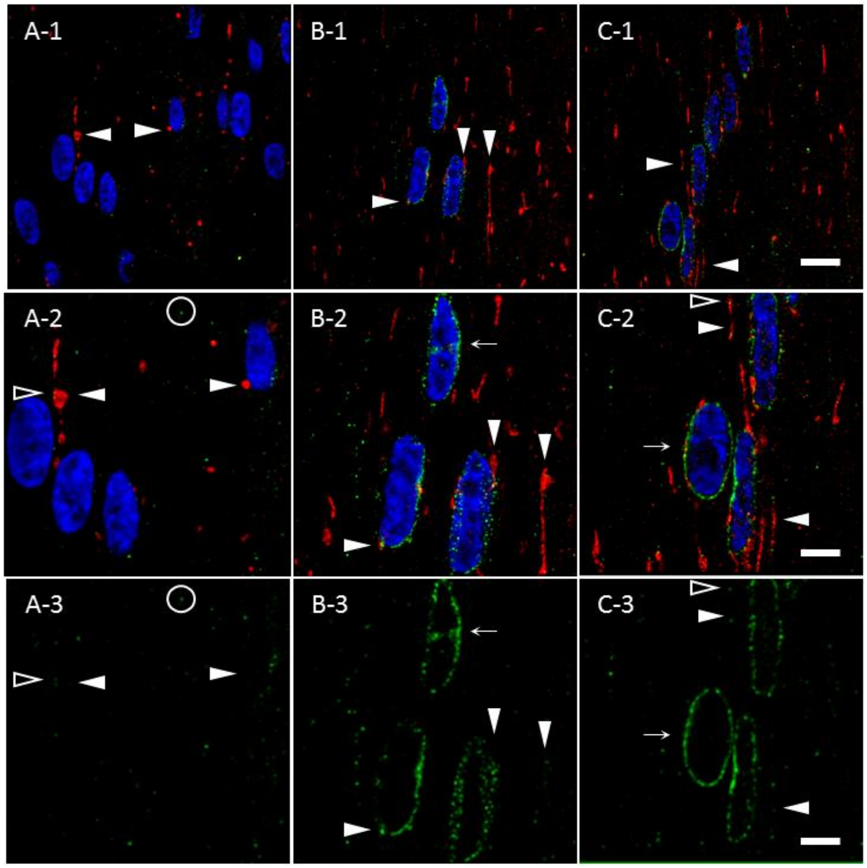


**S4. TOMM20-containing cytoplasmic vesicles lack significant Sec22β expression in bovine lens cortical fiber cells.**

***A-1* – *C-1*.** High-magnification images of *Sec22β* immunolabeling (***green***) and *TOMM20* immunolabeling (***red***) in the *peripheral outer cortex* (**A**), *medial outer cortex* (**B**), and *outer cortex-inner cortex transitional region* (**C**) as demarcated in **Figure 3A**.

TOMM20-containing cytoplasmic vesicles (**closed arrowheads**) are expressed throughout the outer cortex. Initial *Sec22β* expression in the *peripheral outer cortex* (**A**) is weak but strengthens within the *medial outer cortex* (**B**) and remains consistent within the rest of the outer cortex in **C**. Throughout the outer cortex, *Sec22β* is initially expressed within diffuse, punctate cytoplasmic vesicles (**A**, *hollow circle*) that tangibly but insignificantly colocalize with TOMM20-containing cytoplasmic vesicles (***A-2***, ***A-3***, ***C-2***, and ***C-3***; *open arrowheads*). Excluding the lens modiolus, *Sec22β* is also ubiquitously expressed within the cellular nucleus-contiguous rough endoplasmic reticulum (***B-2***, ***B-3***, ***C-2***, and ***C-3***; *solid arrows*).

***A-2* – *C-2*.** Enlarged images of the TOMM20-containing cytoplasmic vesicles demarcated by arrowheads in ***A-1***, ***B-2,*** and ***C-1***.

***A-3* – *C-3*.** Replicate images of ***A-2*** – ***C-2*** depicting *Sec22β* immunolabeling and DAPI labeling only.

Scale bars denote 10 µm (***A-1*** – ***C-1***) and 5 µm (***A-2* – *C-2*** and ***A-3* – *C-3***).
